# KDM4-dependent DNA breaks at active promoters facilitate +1 nucleosome eviction

**DOI:** 10.1101/2023.07.14.548993

**Authors:** László Imre, Péter Nánási, István Szatmári, Endre Kókai, Caroline A. Austin, Viktor Dombrádi, Gábor Szabó

## Abstract

When the effect of various posttranslational histone tail modifications (PTMs) on nucleosome stability was compared in an *in situ* assay involving agarose-embedded nuclei, the promoter proximal H3K4me3, H3K27ac and H4K8ac positive nucleosomes exhibited relative sensitivity to intercalators as compared to bulk H3-GFP or nucleosomes carrying any of the following marks: H3K27me1, H3K27me2, H3K27me3, H3K9me1, H3K9me2, H3K9me3, H3K36me3, H3K4me0, H3K4me1, H3K4me2, H3K9ac, and H3K14ac. Nickase or DNase I treatment of the nuclei, or bleomycin treatment of live cells, did not affect the stability of nucleosomes carrying H3K4me3 or H3K27ac, while those of the second group were all destabilized upon treatment with intercalators. These observations support the possibility that the promoter proximal marks specify dynamic nucleosomes accomodating relaxed DNA sequences due to DNA breaks generated *in vivo*. In line with this interpretation, endogeneous, 3’OH nicks were mapped within the nucleosome free region of promoters controlling genes active in human mononuclear cells, a conclusion supported by superresolution colocalization studies. The +1 nucleosomes were stabilized and the incidence of nicks was decreased at the promoters upon KDM4a,b,c KO induction (Pedersen et al, EMBO J, 2016) in mouse embryonic stem cells (mES). While etoposide did not further destabilize +1 nucleosomes in control mES, their stabilized state in the KO state was reversed by the drug. A significant fraction of the DNA breaks comprises TOP2-generated nicks according to the results of molecular combing experiments. The chromatin regions harboring nicks are topologicaly separated from the domains containing superhelical chromatin. These observations lend support for a model where the role of DNA strand discontinuities in transcriptional regulation and in higher-order chromatin organization are integrated.

## Introduction

DNA breaks have been detected at certain promoters during RNA polymerase II (RNA Pol II) transcription initiation, usually attributed to topoisomerase II (TOP2) cleavages on the grounds of the 3’OH character of the free DNA ends detected upon gene activation and the presence of the enzyme in the protein complex together with the polymerase ^1, 2, 3, 4, 5, 6^. It is hypothesized that the role of these gene activity dependent, transient breaks is to prevent the development of superhelical tension in front of and behind the transcriptional bubble and/or to overrule the topological constraints imposed on enhancer-promoter interactions. The promoter proximal localization of the breaks cannot be readily interpreted in the context of the first, while there is no experimental evidence for the second. The data on gene specific breaks are reminiscent of earlier observations made at a global level, of preformed DNA breaks in the genomic DNA of healthy, nonapoptotic eukaryotic cells including yeasts ^7, 8, 9, 10, 11, 12^. DNA lesions at regulatory regions arising in the course of the generation and repair of histone and DNA demethylation-related oxydative reactions have also been reported ^13^. These breaks as well as the topoisomerase I (TOP1) -related DNA breaks of similar location appear to be under the tight surveillance of a protein complex involving NuMA and members of the single-strand (ss) break repair machinery and are hypothesized to serve the purposes of resolving superhelical tension emerging during looping or the pause-release step of transcriptional elongation ^14^.

Overcoming nucleosomal repression is a fundamental regulatory principle of gene regulation in eukaryotes^15^. For gene activation, the promoter must acquire a nucleosome-free state (leading to nucleosome-free regions, NFRs) in a process aided by remodelling enzymes, histone chaperones ^16, 17^ and TOP1 ^18^. As a result, the negatively superhelical, i.e. underwound, hence melting-prone DNA that forms left-handed toroidal superhelices on the nucleosomes would become interconverted into pectonemic writhes, possibly stabilized by proteins ^19, 20^ until the large RNA POL II preinitiation complex assembles, recognizing promoter specific sequence motives perhaps in the topological context provided by the plectoneme ^19^. Formation of the open complex involves local melting of a short region of the promoter DNA and appropriate positioning of the template strand within the polymerase ^21, 22^. Melting would be facilitated by negative supercoiling ^22, 23^, however, this may not be the case for the plectonemic NFR. The polymerase, following a brief elongation period, is thought to wait for signals to escape from a relatively stable, poised state ^24, 25^, coincident with remodeling of the promoter proximal +1 nucleosome ^17^. The pausing RNA POL II peak is located at + 110 bp relative to the transcription start site (TSS) ^26^. During elongation, the nucleosomes are partially disrupted and get reassembled in the wake of Pol II ^27^. The nucleosomes themselves exhibit remarkable structural plasticity whereby they can accommodate DNA in their different superhelical states ^28^. Stability of the nucleosome is determined by the highly dynamic molecular contacts between histones and the 1.65 helical DNA turns wrapped around their octamer ^29, 30^, and by protein-protein interactions within and, in higher-order structures, between nucleosomes. Destabilization of the nucleosome (octasome, ^31^) may be initiated at the regions where DNA enters and exits the spool, leading first to dissociation of the H2A – H2B dimers yielding hexasomes and eventually tetrasomes ^28, 32^. The elongation coupled over-and underwinding of the DNA double helix, in front of and after the polymerase, respectively, may be concurrent with the formation of these transient structures, suggested also by single molecule experiments when a dramatic loss of H2A/H2B dimers ensues under positive torque ^33, 34^. The tetrasomes, with their right-handed configuration, could absorb the positive extra supercoils generated ahead of the polymerase, and revert to lefthanded chirality upon rebinding of the H2 dimers, behind ^34, 35^. These effects may spread far beyond the point of actual transcription ^23, 36^. During transcription, superhelicity is effectively regulated by topoisomerases that can change the number of intertwinings of the two DNA strands in cleavage-religation reactions working in collaboration with chromatin remodellers ^18 37^. The presence of transient DNA breaks cannot be readily fit in the above complex picture.

The histone tails that protrude from the nucleosomes and their posttranslational modifications (PTMs) may play important but partly understood roles in the regulation of this elaborate machinery. The active mark H3K4me3 e.g., recruits the Mediator complex ^38, 39^ and the chromatin remodeling protein CHD1 to upregulate nucleosome turnover at most Pol II-directed promoters, to allow efficient polymerase escape ^17, 39^. The nucleosome-DNA binding is stabilized by the tails that contact the internucleosomal linker DNA ^40^ and also mediate internucleosomal interactions ^41^. Wedging into the minor groove, the tails gate the access of the polymerase ^42, 43^ and may also control the tetrasome chiral transition accompanying elongation ^44^ ^45^. Studies in cell free systems raise the possibility that the tails themselves also have an autonomous influence on nucleosome stability, especially in the case of tetrasomes ^46, 47^. Proteolytic removal of the N-termini of the core histones leads to functional consequences comparable to histone acetylation, suggesting that this PTM destabilizes nucleosomes by reducing the stability of interactions mediated by unmodified tails ^48^. Alternatively, nucleosome stability might be modulated by reader proteins recruited by the tails, in an unknown manner. The possible regulatory effect of superhelicity relaxation on nucleosome stability has not been throroughly investigated before, to our knowledge.

To further investigate these complex relationships, we have developed an *in situ* assay for the analysis of the effect of the different histone modifications on nucleosome stability on a cell-by-cell basis. Through the spectacles of this approach, nucleosomes flanking the nucleosome free regions (NFRs) appear to be specifically destabilized at active promoters and the mechanism of destabilization involves DNA relaxation ensuing in the wake of DNA break formation there. Through the spectacles of our *in situ* approach ^49^, the superhelical state of the DNA emerges as a primary determinant of nucleosome stability. Both the stability of the +1 nucleosomes and the free 3’OH DNA breaks in their vicinity was KDM4-dependent and appeared to include topoisomerase-generated single-strand (ss) breaks, nicks. The data presented reveal that nucleosome stability, histone PTMs and DNA topology are interconnected determinants of transcriptional regulation involving the +1 nucleosomes. We also show that the nicks are topologically separated from their superhelical chromatin environment.

## Results

Exploiting the advantages of laser scanning cytometry, elution profiles of bulk nucleosomal histones and their different posttranslationally modified species were recorded on a cell-by-cell basis. (see brief description of the method (ref. ^49^) in Suppl. Fig. 1). When the agarose embedded nuclei were exposed to increasing concentrations of intercalator solutions, the amount of different histones remaining in the individual nuclei rapidly declined above a certain concentration of the eluent (Fig. 1, A). The destabilizing effect of the intercalator EBr was observed only in the presence of at least 750 mM NaCl in the case of H3 or H4, as described in ref. ^49^ Nucleosomes decorated with the active marks H3K4me3, H3K27ac and H4K8ac were markedly less stable then those marked by H3K27me1, H3K27me2, H3K27me3, H3K36me3, H3K9me1, H3K9me2, H3K4me0, H3K4me1, H3K4me2, H3K9ac, H3K14ac and H4K16ac (Fig. 1A-F), bulk H3-GFP or H4-GFP containing nucleosomes (Suppl. Fig. 2A, B). Using the clinically relevant intercalator Doxorubicin, the pattern of PTM dependent sensitivity was identical to what was obtained with EBr (Suppl. Fig. 2C, D). Doxorubicin destabilized nucleosomes also at low salt (150 mM NaCl).

**Fig. 1.**
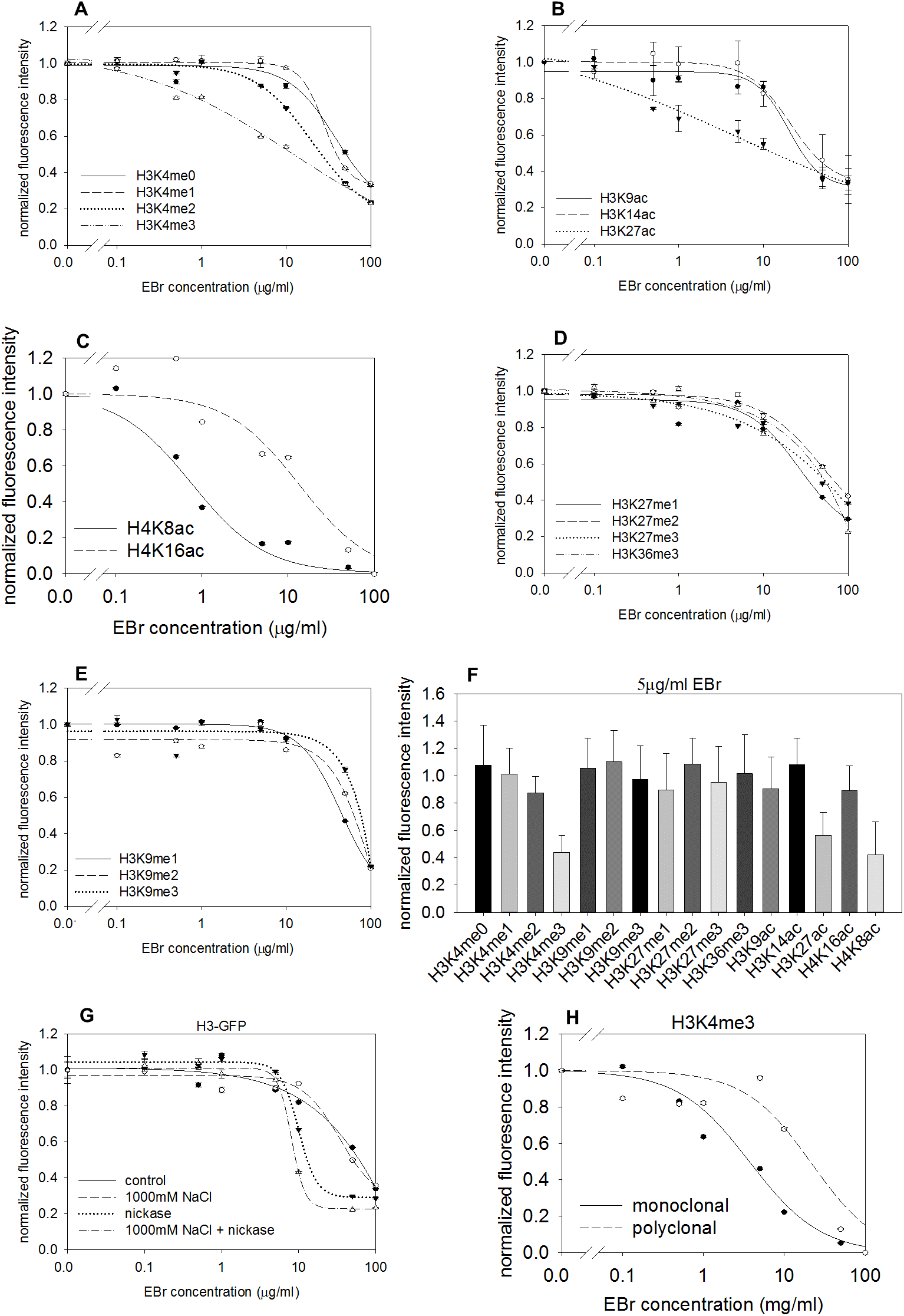
Effect of histone PTMs, partial nucleosome depletion and DNA relaxation on nucleosome stability. (A-E) A panel of antibodies specific for histone modifications characteristic for euchromatin: H3K4me0/1/2/3 (A), H3K9/14/27ac (B), H4K8/16ac (C), H3K36me3 (D) or heterochromatin: H3K27me1/2/3 (D), H3K9me1/2/3 (E) was used to detect the amount of the different PTM-marked nucleosomes in the QINESIn assay ^49^, in H3-GFP or H4-GFP expressor HeLa nuclei. The intercalator ethidium bromide (EBr) was applied as nucleosome destabilizing agent and a panel of PTM-specific antibodies ^84^ for the characterization of the stability of different nucleosomes. H3-GFP and H4-GFP were used as internal control (Suppl. Fig. 1A and B). (F) Comparison of nucleosomes marked by different PTMs using 5 µg/ml EBr (and 750 mM salt; see ref. ^49^) in the case of HeLa nuclei. (G) EBr elution assay of H3- GFP expressing HeLa nuclei after nucleosome depletion by pretreatment with 1M NaCl and after topological relaxation by a nickase enzyme. Initial H3-GFP fluorescence intensities in the absence of EBr are shown in Suppl. Fig. 1F. (H) Comparison of H3K4me3 specific monoclonal ^84^ and polyclonal (Abcam, ab8580) antibodies in the EBr elution assay, in H3-GFP expressor HeLa nuclei. H3-GFP was used as an internal control, as shown in Suppl. Fig. 1E. The curves refer to 200-2000 G1 phase cells gated according to their DNA content. Bars represent SEM.

Since H3K4me3, H3K27ac, and also H4K8ac ^50, 51, 52^ are attributes of the nucleosomes flanking transcriptional start sites of active promoters, our working hypothesis was that the +1 nucleosomes are destabilized relative to nucleosomes outside these chromatin regions. Decreased stability of the +1 nucleosomes may not be the result of the presence of nucleosome-free regions in their vicinity, since depletion by ∼30% of the nucleosomes prior to intercalator elution did not change bulk H3-GFP stability (Fig. 1G, Suppl. Fig. 2E). On the other hand, relaxation of internucleosomal superhelicity by nickase treatment elicited significant nucleosome destabilization in both the control and salt-pretreated nuclear samples (Fig. 1G).

Interestingly, the H3K4me3 nucleosomes exhibited destabilized character only when detected by certain H3K4me3-specific antibodies (Fig. 1H; see inner control in Suppl. Fig. 2F, suggesting that they recognize or access different fractions of the nucleosomes carryining these epitopes, what may not be reflected by ChIP-seq maps obtained by the different antibodies, in view of the fundamental dissimilarities in the antibody labeling protocol of the ChIP-seq and elution experiments. (If not specified, the monoclonal antibody detecting unstable H3K4me3 nucleosomes was used in the experiments.)

As opposed to the sensitivity of bulk nucleosomes or inactive mark labeled nucleosomes (Fig. 2 A, Suppl. Fig. 2G, H) to nicking, the same treatment did not affect the stability of nucleosomes carrying H3K4me3 or H3K27ac (Fig. 2B). The disparate effect of nicking, as measured in the case of HeLa nuclei, was reproduced also in the case of lymphocytes isolated from human peripheral blood (hPBMCs; Fig. 2C, D, Suppl. Fig. 2I, J), and mouse embryonic stem cells (mES; see control samples of Fig. 5A-D, below). The stable H3K4me3 nucleosomes identified by the polyclonal antibody were also sensitive to nickase treatment (Suppl. Fig. 2K). Nicking the DNA within live cells using bleomycin, an agent known to generate ss breaks *in vivo*, also destabilized H3-GFP and H3K27me3, while H3K4me3 and H3K27ac were not affected (Fig. 2E, Suppl. Fig. 2L, M). Treatment with DNase I also leads to destabilization of bulk H3 both when used in conditions favoring single-strand cleavages and when applied at high concentration, without significantly affecting the stability features of the H3K4me3 nucleosomes (Fig. 2F, Suppl. Fig. 2N, O).

**Fig. 2.**
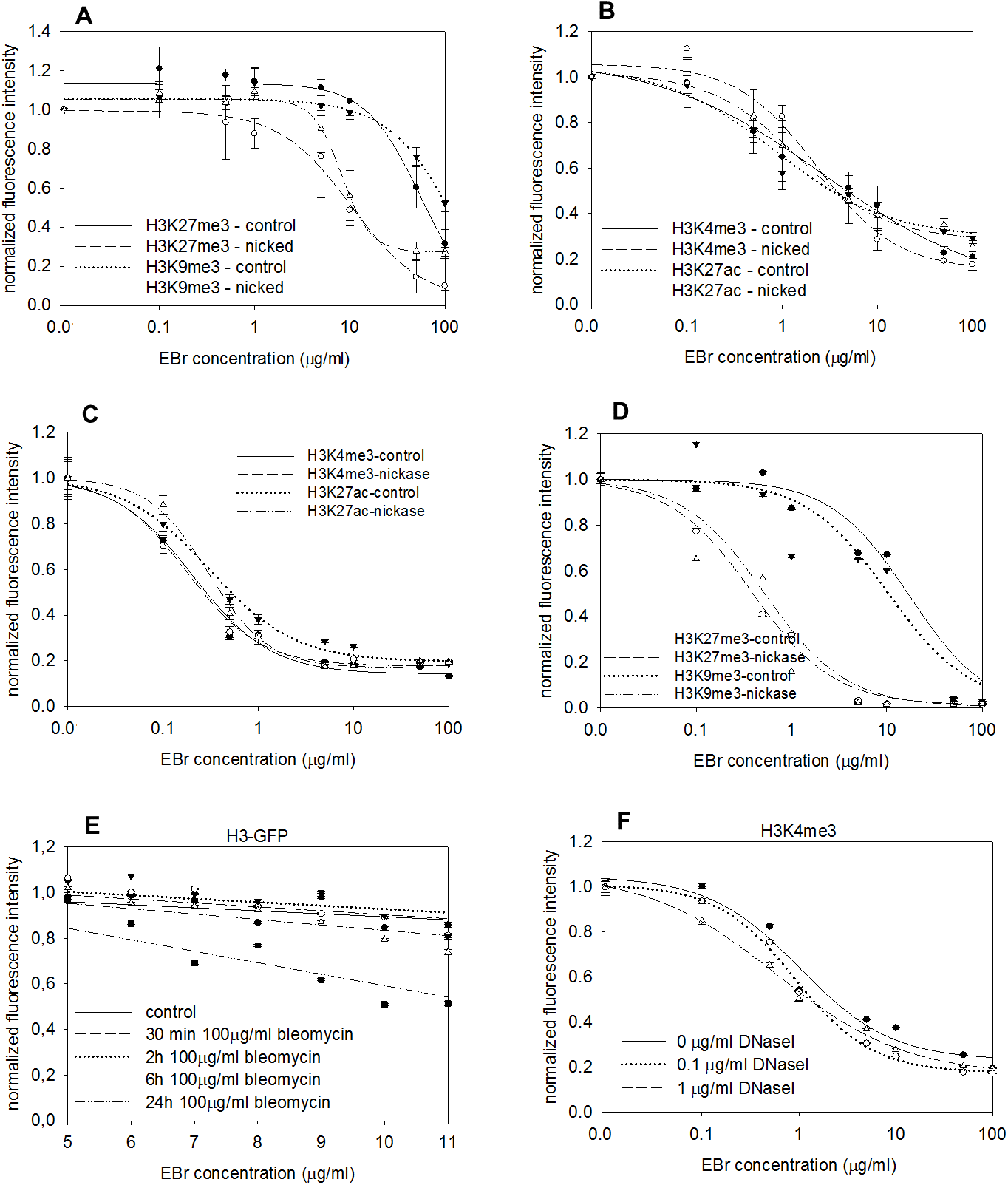
Effect of DNA relaxation on the stability of nucleosomes at the TSS or in heterochromatin. (A and B) EBr elution measured in H3-GFP expressor HeLa cell nuclei. The nucleosomes remaining in the nucleus at the different intercalator concentrations were detected by H3K27me3, H3K9me3 (A) or by H3K4me3, H3K27ac (B) specific antibodies, following relaxation by nickase treatment, or in its absence, as indicated in the figures. H3-GFP was used as an inner control as shown in Suppl. Fig. 2G, H. (C and D) EBr elution of H3K4me3, H3K27ac, H3K27me3 or H3K9me3 marked nucleosomes in relaxed (nickase) or superhelical (control) chromatin of HPBMC nuclei. Elution curves measured in the nuclei of cells isolated from another healthy donor are shown in Suppl. Fig. 2I, J. (E) EBr elution curves measured in the nuclei of H3-GFP expressor HeLa cells treated with bleomycin for different times. (F) EBr elution curves measured in H3-GFP expressor HeLa cell nuclei digested by different concentrations of DNase I before elution and labeling of H3K4me3. Elution curves of H3-GFP used as inner control are shown on Suppl. Fig. 2N. Initial fluorescence intensities at no EBr are shown in Suppl. Fig. 2O. The curves refer to 200-2000 G1 phase cells gated according to their DNA content. Bars represent SEM.

The peculiar lack of any effect of nicking on the stability of the H3K4me3 and H3K27ac decorated nucleosomes raised the possibility that these marks specify dynamic nucleosomes accommodating *already relaxed* DNA sequences, while most other nucleosomes hold the DNA in constrained superhelices. In line with this possibility, using nick-labeling and DNA immunoprecipitation (NLDI; see Materials and Methods), 3’OH nicks were mapped to the vicinity of gene promoters of the G0 human peripheral blood cells, of active genes in particular (Fig. 3A, B), in line with some of the data obtained by altenative strategies of break-mapping ^53^. Intriguingly, the lists of promoters harboring nicks were overlapping to a significant degree (Fig. 3C, D and Suppl Fig. 3), suggesting that a considerable fraction of the nicks detected is constitutive. Images obtained by superresolution microscopy (STED) show co-localization of the nicks labeled by nick-translation and RNA POL II, especially in the case of its Ser 5-phosphorylated initiating form (Fig. 3E-I).

**Fig. 3.**
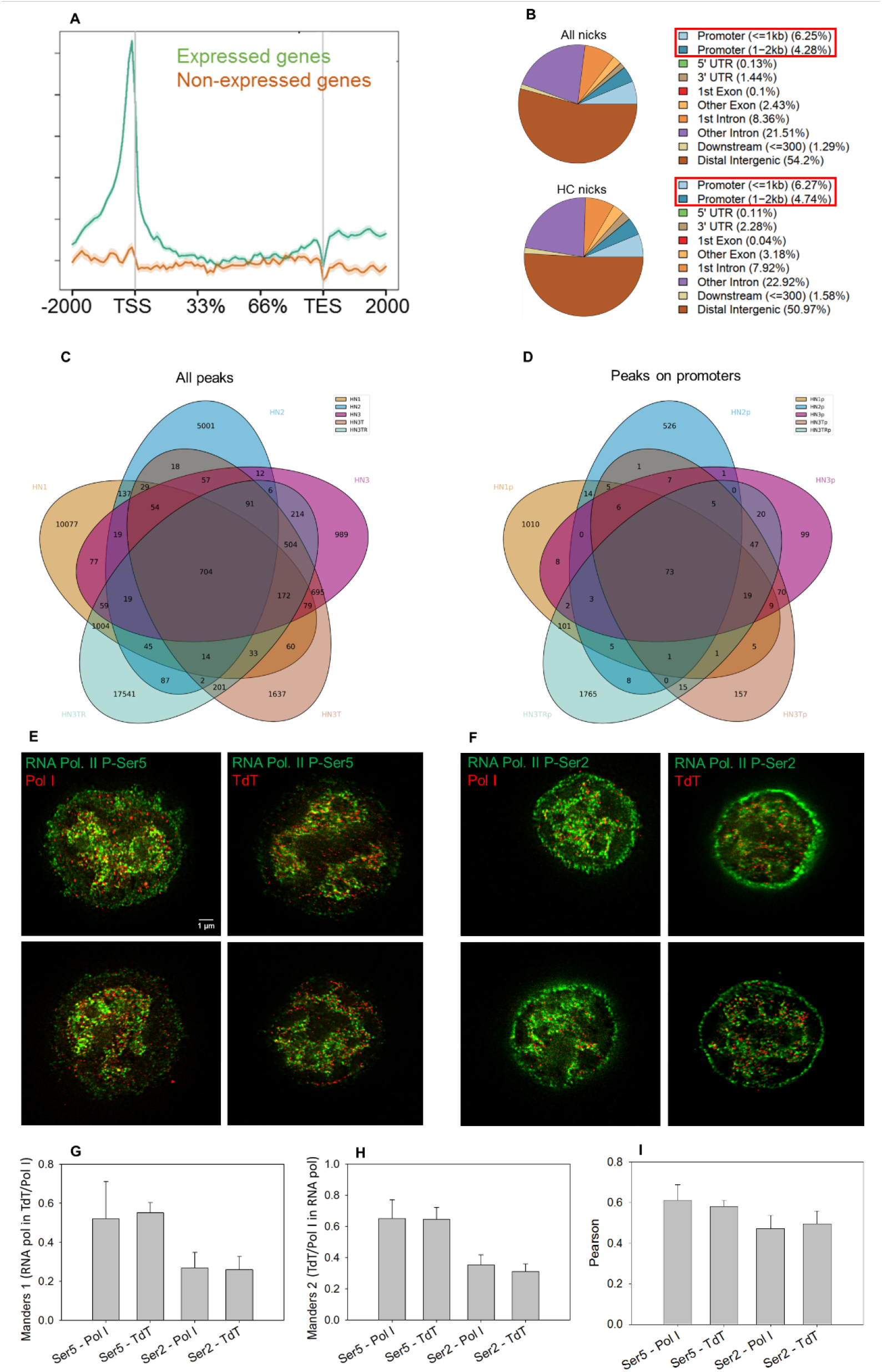
Distribution of 3’ OH breaks in hPBMC DNA. (A and B) Biotin-labeled nucleotides were incorporated into agarose-embedded deproteinized hPBMC DNA using E coli DNA polymerase I holoenzyme (DNA POL. I), followed by immunoprecipitation and sequencing (NLDI; see Materials and Methods). Panel A shows the metaplots of active and inactive genes (annotated based on their RNA-seq FPKM values, as described in Materials and Methods), shown by the green and brown line, respectively. The Y axis shows read count per million mapped reads, the X axis represents the genomic region. Panel B shows the annotation of NLDI peaks by ChIP-seeker for peaks detected in all 4 and in at least 2 independent experiments („All nicks” and „HC nicks”, respectively). ∼10% of peaks localize to promoter proximal (+/- 2kb) genomic regions. (C and B) Venn-diagrams comparing peaksets from 5 independent nick-labeling experiments. All peaks (left) and peaks detected on promoters (right) are compared. „HN1-3” represents peaksets in the case of nick-labeling by DNA POL I, „HN3T” represents nicks labeled by TdT, while in „HN3TR” nicks were labeled by TdT followed by RNAse A treatment to remove any possible signal of RNA end-labeling. Peaksets ending with „p” show promoter peaks. (E-I) DNA breaks partially co-localize with RNA POLymerase during transcription initiation and elongation. STED microscopic images of agarose-embedded, fixed Jurkat cells. DNA breaks were labeled by DNA POL I or TdT, the Ser5 phosphorylated (E) and Ser2 phosphorylated (F) forms of RNA POL II were visualized using antibodies specific for the differently phosphorylated forms of the enzyme. The incorporated biotin was detected by streptavidin conjugated Abberior Star Red (red). Panels G and H show co-localization of DNA breaks with Ser5 and Ser2 phosphorylated RNA POL II, respectively. (G) Distribution of co-localization values (Pearson and Manders 1 and 2) calculated from the samples of (E) and (F). The error bars represent SEM values. (5-5 STED microscopic images were used for quantification per sample). Representative STED microscopy images are shown in (E) and (F).

The 3’OH DNA breaks detected are likely ss breaks, since our labeling strategy involved nick-labeling using DNA polymerase I (DNA POL I). This was confirmed by incorporating biotin-labeled nucleotides into deproteinized genomic DNA of agarose-embedded HPB cells, when DNA was isolated and combed onto coverslips and the nick signals were observed mainly within the combed, loop-size fragments, rather than at the ends (Suppl. Fig. 4A-F). The majority of the labeled nicks could be co-labeled using biotin-conjugated recombinant TDP2, suggesting that they carried proteolytic remnants of TOP2 enzyme attached covalently to the 5’-P of the nick (Suppl. Fig. 4G-I). The identity of the topoisomerase enzyme responsible for the breaks was likely TOP2β, rather than TOP2α, in view of the G0 character of the hPBMCs used in these experiments. In line with this assumption, significantly decreased levels of nick-labeling were found in 2 TOP2β -KO cell line, as shown in Suppl. Fig. 5.

We have mapped endogenous ss breaks by NLDI also in mouse embryonic stem cells, before and after induction of knocking out of three KDM4 genes (KDM4a,b,c; ^54^; experimental system kindly provided by C. Helin and K. Agger). The incidence of promoter-proximal nicks was decreased upon KDM4 KO (Fig. 4A- D), a change deemed significant in view of the large background of Okazaki fragment related DNA breaks in these proliferating cells. Using the KDM4 KO mouse embryonic stem cell model we investigated if the +1 nucleosomes, recognized based on their H3K4me3 content become stabilized through the spectacles of intercalator elution. As Fig. 4E-J shows, there was a considerable and reproducible shift in the stability of H3K4me3 nucleosomes, as opposed to that of the H3K9me3-marked heterochromatic nucleosomes (Suppl. Fig. 6). Relaxation by nickase (Fig. 5A-D), or treatment with etoposide (Fig. 5E-J, Suppl. Fig. 7) shifted back the stability of the H3K4me3 nucleosomes to those of the control, showing that topoisomerase II was in place and potentially active, but rendered inactive in the absence of KDM4 or without the enzymatic activities involved. The decrease of nicks upon KO induction was also analyzed in an experiment when the overall nick levels were studied by a reverse Southwestern (rSW) blot procedure ^55^, as shown in Suppl. Fig. 8. The overall decrease was about 10 %, as compared with the ∼50% reduction of promoter-proximal nicks in the DIP-seq experiment, suggesting that the latter class of DNA breaks were selectively affected by KDM4 KO.

**Fig. 4.**
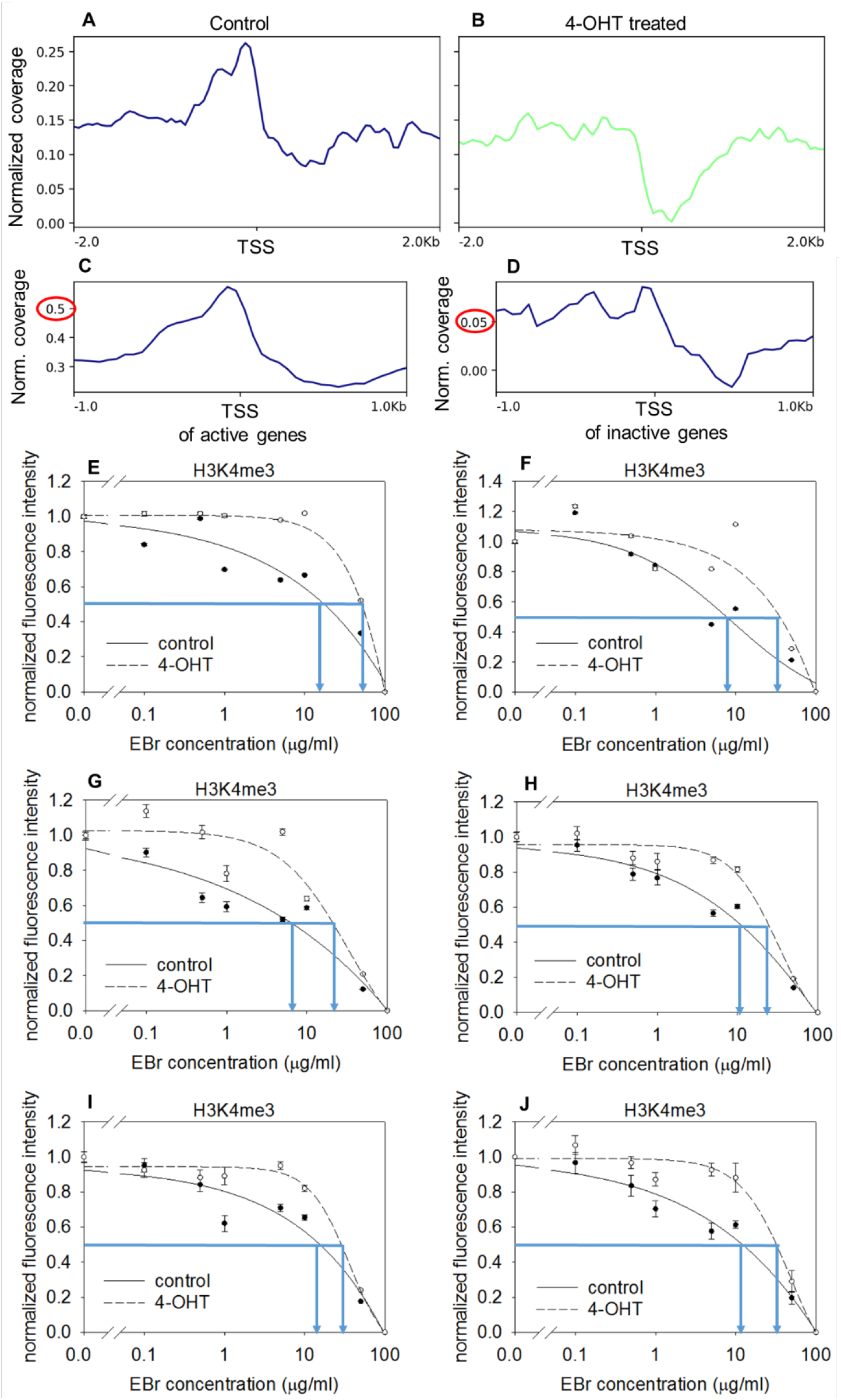
KDM4-dependent breaks detected the TSSs of mES cells and nucleosome stability measured in the nuclei of 4-OHT inducible KDM4a,b,c KO mES cells. (A and B) Distribution of 3’ OH DNA breaks detected by NLDI in mES cells before (A) and after 4-OHT (B) treatment. Read coverages were normalized to input control. (C and D) Anchor plots show the normalized nick coverage around the TSSs of active (C) and inactive (D) genes. Expression was determined based on microarray data (GSE64251; ^54^. Genes were diveded into active and inactive categories based on their FPKM values. (E-J) EBr elution curves of H3K4me3 nucleosomes measured in control or 4-OHT induced KDM4 KO mES nuclei. Results of independent experiments are shown using H3K4me3 specific IgG (E-G) or, to increase accessibility, its Fab fragment (H and I). The curves refer to 200-2000 G1 phase cells gated according to their DNA content. Bars represent SEM. Average values and SEM of the five independent measurements are shown in panel (J). Elution curves representing H3K9me3 nucleosomes co-labeled with H3K4me3 are shown in Suppl. Fig. 6.

If the destabilized state of the +1 nucleosomes is primarily the consequence of a topologically relaxed state of the DNA region involved, they must be topologically isolated from the neighboring, unrelaxed chromatin regions. We first investigated how long stretches of chromatin are affected by the topological relaxation induced by a nickase enzyme. The concentration of nickase necessary for inducing near-complete H2B-GFP or H3-GFP eviction was determined (see Fig. 6A), and the average distance between the nicks was assessed by treating the chromatin with S1 nuclease followed by non-denaturing agarose gel electrophoresis (Fig. 6B). These studies have revealed that topological relaxation by inducing nicks as rarely as one nick per >10 kb DNA can lead to destabilization of the nucleosomes in the entire chromatin loop. Thus, accumulation of breaks within the nucleosome free regions of promoters would also be expected to relax the neighboring chromatin regions marked with other PTMs (what was not seen; see Fig. 1), unless they are topologically isolated. To investigate if the promoter regions encompassing the +1 nucleosomes indeed form distinct topological domains insulated from the gene regions of transcriptional elongation, nuclear halo and loop-winding experiments were performed using Jurkat cells found optimal for such studies ^49^. The DNA loops protruding out from the lamina-bounded nucleus could be relaxed and then overwound by adding increasing concentrations of an intercalator (Fig. 6C), what is an evidence of the superhelicity of the DNA in the halo ^12, 56^. Nicks could be detected close to the lamina, at the stem of the protruding, superhelical loops (Fig. 6D- G). Thus, genomic DNA comprises two distinct topological domains: superhelical chromatin loops and relaxed regions associated nicks that also colocalize to a significant degree with the +1 nucleosomes and appear to be anchored to the lamina.

## Discussion

We have observed a major difference between the stability of nucleosomes marked with H3K4me3 and H3K27ac as well as H4K8ac, as compared with all other nucleosomes tested (Fig. 1). H3K4me3 is generally considered as a hallmark for the promoter proximal +1 nucleosome of active genes ^57^ and refs. cited therein, while H3K27ac is deposited on the same nucleosomes and also on those flanking active enhancer elements^58^. H4K8ac was also mapped near the TSS of active genes ^51, 52^.

The lack of an effect of nicking in the case of these H3K4me3, H3K27ac or H4k8ac marked nucleosomes is best interpretated assuming that the DNA here is already relaxed. Intercalating into either intact or nicked DNA, EBr (similarly to Sybr Gold and Doxorubicin) is expected to change the twisting of the double helix^59^, hence stretching the DNA, what may lead to displacement of the contact sites between nucleosome and DNA, so the nucleosomal barrier mounting before transcription can be overcome.

At the concentrations eliciting eviction (Fig. 1), intercalators, decrease the negative superhelical twist within the nucleosomes, cause further extension relative to their baseline stretching and compete with the core histone tails at the outskirts of the nucleosomes, in line with their demonstrated interactions with linker DNA ^60, 61, 62, 40^. The N-terminal H3 tails significantly contribute to the superhelical constraint imposed by the tetrasomes on the DNA ^45, 63^, therefore competition with H3 histone tails what may involve most nucleosomes but the H3K4me3 marked ones, could explain why the latter nucleosomes are selectively unaffected by nicking. However, elution by EBr occurs only in the presence of at least 750 mM salt; at this salt concentration the tails do not interact with the DNA and there are no internucleosomal interactions either ^64^, as the salt-bridge interactions involved are disrupted ^65^. Nicking *in vivo* by bleomycin destabilized bulk nucleosomes only (Fig. 2), suggesting that relaxation induced nucleosome destabilization may occur also in live cells. Indeed, we have mapped *bona fide* nicks at active promoters (Fig. 3). These mapping experiments involved cells fixed in 1 % formaldehyde, suggesting that the nicks exist *in vivo*.

The torsional state of DNA *in vivo* is assumed to be determined primarily by its nucleosomal structure ^66^ and it is dominated by the negative superhelical twist constrained by several histone-DNA contacts ^30^. Negative superhelicity is thought to assist transcriptional and replicational processes by facilitating melting of the double-helix, what is one of the rate limiting steps in transcriptional initiation, besides preinitiation complex (PIC) assembly and polymerase pausing ^21^. Due to its constrained nature, as well as the fact that PIC assembly occurs in nucleosome depleted regions where superhelicity is manifest in plectonemic writhes ^19^, it is not obvious if and how negative superhelicity becomes of advantage for transcriptional initiation. On the other hand, the role of negative superhelicity may involve a further aspect as well, related to its association with nucleosomal structure. We suggest that the H3K4me3-marked, promoter proximal, highly positioned nucleosome +1 contains DNA in a relaxed state, decreasing nucleosomal stability. Since the RNA POL II must overcome the thermodynamic barrier imposed by the first nucleosome so as to become released from a poised/paused state ^17, 24, 26, 67, 68^, relaxation of the DNA in this region may have regulatory significance. Transient nicks, though generated by TOP I, are associated with eRNA synthesis and enhancer activation according to a recent report ^69^, widening the scenario where relaxation evoked nucleosome destabilization might play a role. Although TOP1 also binds at the gene promoters ^18^, the 3’ ends would be unavailable for labeling by Klenow, hence go undetected in our mapping experiments. The most likely mechanism of relaxation is nicking by TOP2β, since this enzyme was implicated in the generation of the transient strand breaks observed upon activation of certain genes ^2, 4, 45, 70^, and nicking appears to be its dominant mode of its action *in vivo* ^71^. The high degree of microscopic colocalization between nicks and TOP2β (Fig. 6), is in line with this model.

Nucleosome eviction elicited by doxorubicin and other intercalating drugs as a consequence of torsion-induced nucleosome destabilization ^72, 73^ can lead to DNA regions exposed to reactive oxygene species (ROS) and mechanical stress, what can explain the reported DNA breakages observed at promoters of active genes in the wake of anthracycline treatment of cells ^74^. These breaks were mapped to the nucleosomes bordering the nucleosome depleted regions in both yeast and mouse, in line with our suggestion that the endogeneous nicks arise within the DNA wound around the +1 nucleosome. Nicked sites are prone to mechanical breakage so as to form double-strand discontinuities ^55, 75^. We speculate, sharing the views cited above, that they may represent hotspots for recombinational events leading to translocations.

The observation that some of the KDM4 genes knocked out in our experiments (KDM4a,b,c; ^54^) regulate the incidence of 3’OH nicks at promoters as well as the stability of the +1 nucleosomes could be explained by ROS-generation upon histone demethylation here, in the context proposed in ^13^. However, the etoposide-induced reversal of the stability of the KDM4 KO-stabilized H3K4me3 nucleosomes suggest the direct involvement of TOP2 in generating the nicks. In line with this hypothesis, there is a large overlap between the lists of promoters carrying nicks in our experiments and in the case of the TOP2-related DNA breaks revealed in another cell line in the presence of etoposide ^76^, as shown in Suppl. Fig. 9. In a recent study KDM4B was found among the chromatin regulatory genes determining anthracycline sensitivity ^77^ and accessibility of TOP2 to chromatin was proposed to be the basis of sensitization. The etoposide-induced destabilization of +1 nucleosomes in the KDM4 KO mES (Fig. 5) suggests that the enzymatic activity rather than access to chromatin may be KDM4-dependent.

**Fig. 5.**
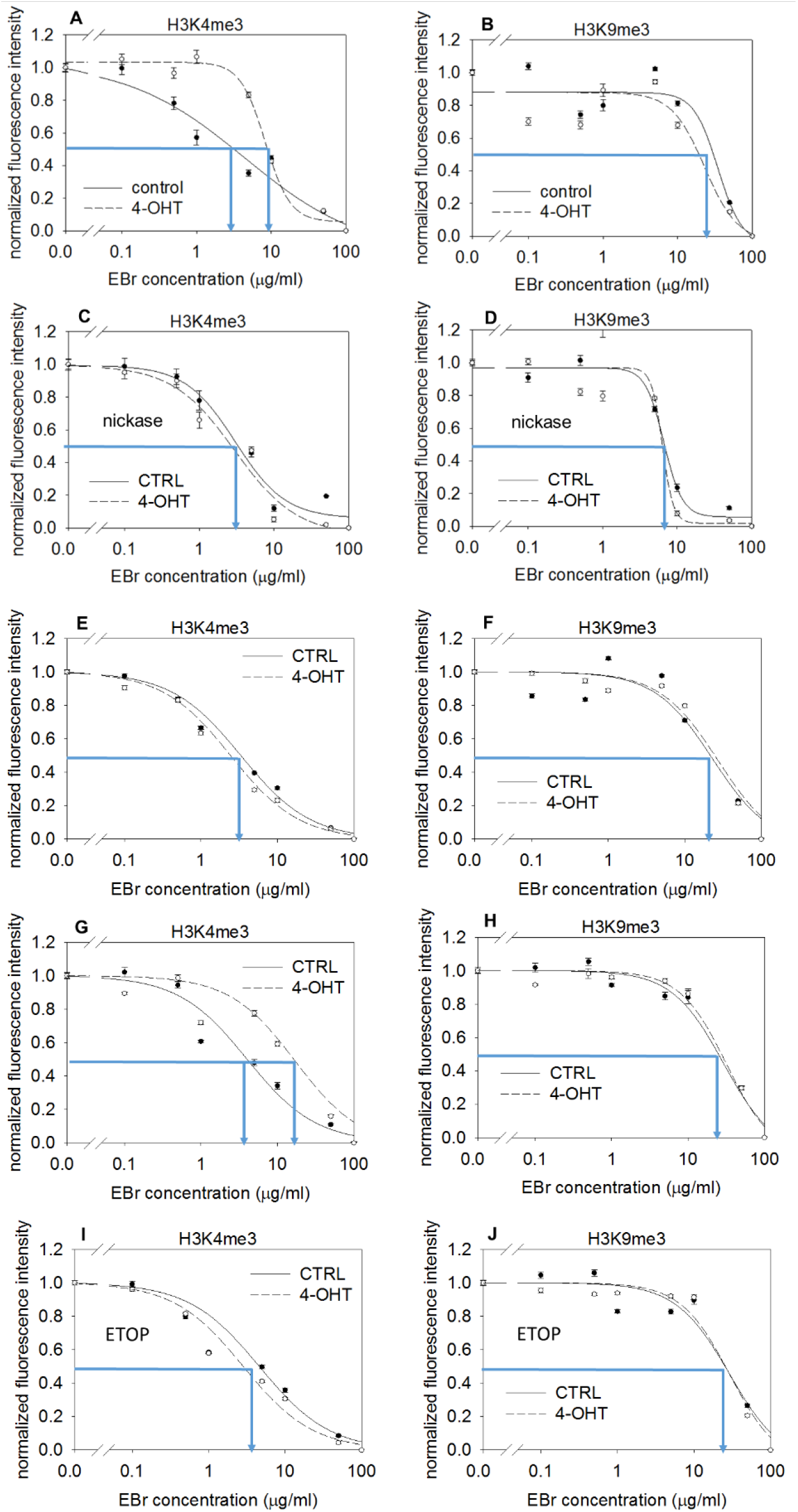
Effect of DNA relaxation by nickase or etoposide treatment on nucleosome stability, measured in 4- OHT inducible KDM4 KO mES nuclei. (A-D) EBr elution curves of H3K4me3 and H3K9me3 nucleosomes measured in control and 4-OHT induced KDM4 KO mES nuclei, treated with nickase (C and D) or in the absence of treatment (A and B). (E-J) EBr elution curves of H3K4me3 and H3K9me3 nucleosomes measured in the nuclei of mES cells lacking the inducible KO construct (E and F). in etoposide (ETOP) untreated mES cells with Jmjd2 KO system (G and H) or in ETOP treated mES cells with Jmjd2 KO system (I and J). Measurements in panels G-J were reproduced in independent experiments shown in Suppl. Fig. 7. The curves refer to 200-2000 G1 phase cells gated according to their DNA content. Bars represent SEM.

Recently, KDM5-dependent H3K4me3 turnover was reported to regulate the recruitment of the integrator complex subunit 11, a step shown to be essential for the eviction of paused RNA POL II ^78^. Perhaps these steps of transcriptional activation follow the H3K4me3 nucleosome destabilization by KDM4/TOP2β- dependent topological relaxation reported herein.

Our data appear to be at variance with studies based on psoralen binding ^79^ suggestive of superhelical strain being maintained at active promoters. However, the increased presence of covalently bound psoralen in these regions may be a consequence of a changing nucleosome landscape in view of the almost exclusive binding of psoralen and other intercalators to nucleosome free and linker regions ^66, 80^.

The colocalization of nicks with RNA POL II molecules also in their elongating state (Fig. 3) may be visualized considering a model where the transcribed DNA loop is reeled through separate cogwheels of initiating and elongating polymarases of the transcriptional factories. In line with this model, elongation and initiation factories do not appear to coincide according detailed superresolution microscopic analyses ^81^. The regions harboring DNA breaks are topologically isolated from those of the superhelical loops (Fig. 6) where transcriptional elongation induces positive and negative changes in internucleosomal superhelicity, in front of and behind the polymerase enzyme, respectively ^82^. This topological separation does not appear to involve steric separation in the nucleus.

**Fig. 6.**
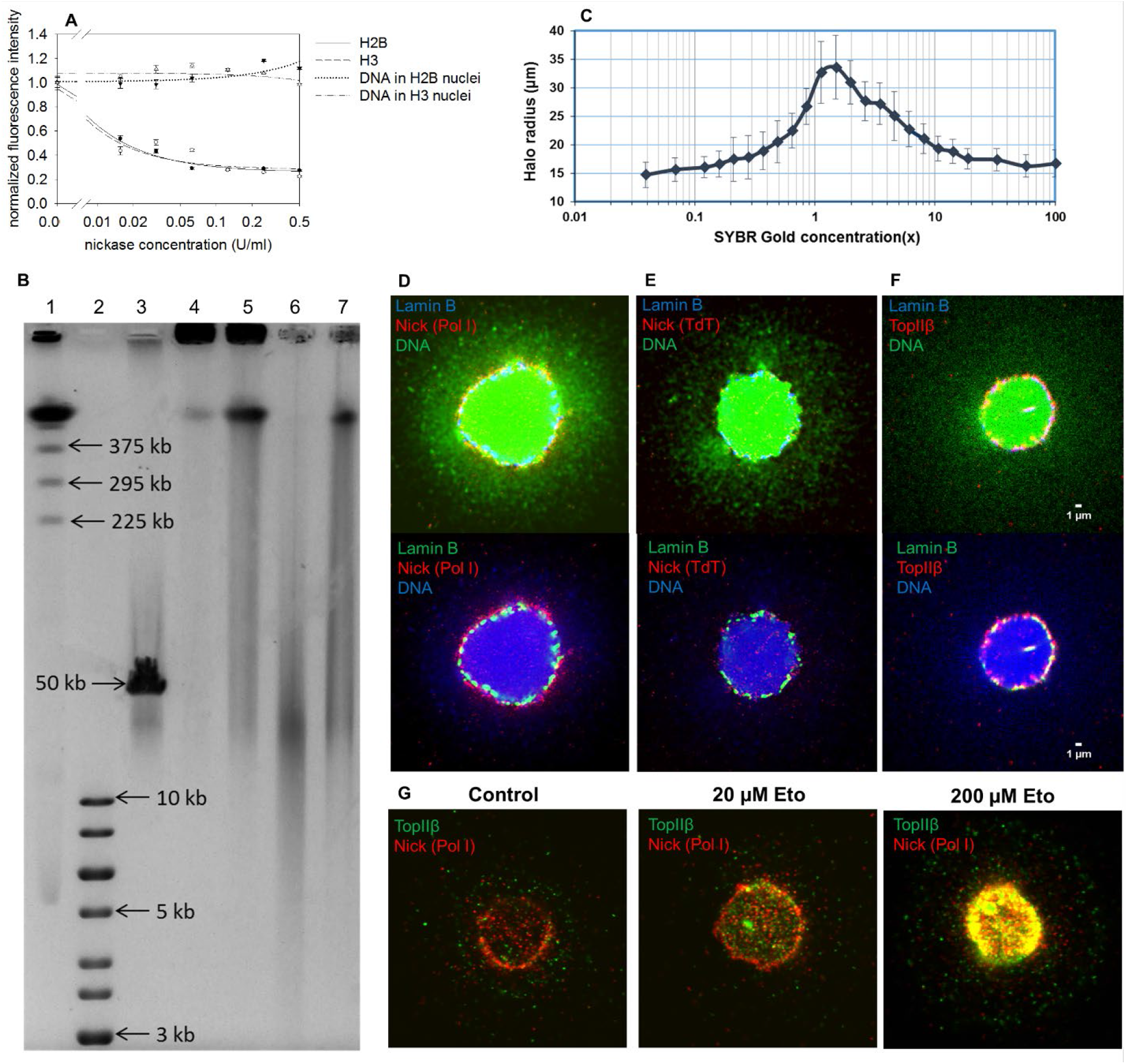
Nucleosome destabilization and nick distribution after nickase digestion. Determination of the superhelical density of eukaryotic chromatin loops. Nuclear localization of nicks. (A) Eviction of H2B-GFP and H3-GFP in nuclei treated with different concentrations of nickase. H2B-GFP and H3-GFP expressor HeLa nuclei treated with 0.95 M NaCl, or 10 µg/ml EBr / 0.75 M NaCl, respectively, after treatment with nickase used at the concentrations indicated in the figure. The curves refer to 200-2000 G1 phase cells gated according to their DNA content. Bars represent SEM. (B) Double-strand fragment size distribution of the DNA after nickase+S1 treatment. The nicks were converted to double-strand breaks by S1 nuclease and the ds fragments analyzed by CHEF as descibed in Methods and Methods. Lane 1: 225-2200 kb Pulse marker; lane 2: 1 kb DNA ladder; lane 3: lambda DNA; lane 4: untreated control; lane 5: 0.5 U/ml nickase; lane 6: 0.5 U/ml nickase + 1000 U/ml S1; lane 7: 1000 U/ml S1. (C) The fluorescent halo radius, determined by laser scanning cytometry, is plotted as a function of SG concentration. Upon addition of the dye the halo radius gradually increases as the negatively supercoiled plectonemic DNA loops progressively uncoil and become increasingly relaxed. Complete relaxation of the loops is observed between 1-1.7X SG concentration. Additional intercalation causes the relaxed DNA loops to form positive plectonemic writhes, thus decreasing the halo radius. (D-E) Nicks detected in nuclear halo samples of Jurkat cells by STED microscopy. Lamin B was detected by immunofluorescence (blue and green in the upper and lower images, respectively), DNA was stained by PI (green and blue in the upper and lower images respectively). Lamin B and nicks (red) only were imaged at superresolution. (F) Lamina-proximal TOP2β was detected on nuclear halo samples by confocal microscopy. Lamin B was detected by immunofluorescence (as before), TOP2β was detected by an antibody prepared in C. Austin’s lab and DNA was stained by PI (as before). The signal in the DNA channel of the upper images of Panels D-F were amplified to better visualize the nuclear halos. (G) Co-localization of lamina-proximal nicks and TOP2β in nulcear halo samples before and after etoposide treatment, visualized by confocal microscopy. TOP2β was detected using a rabbit polyclonal antibody (Santa Cruz: H-286; green)), DNA POL I labeled nicks were detected by anti-biotin antibody (red). Representative microscopic images are shown.

In summary, making use of an *in situ* assay of nucleosome stability, H3K4me3, H3K27ac and H4K8ac PTMs entail striking instability relative to most other nucleosomes. The dynamic character of these nucleosomes could be linked to the relaxed state of DNA due to nicks generated at the active promoters, possibly by TOP2β, and both features are KDM4-dependent. The chromatin regions harboring nicks are topologicaly separated from the domains containing superhelical chromatin. These observations lend support for a model where the role of DNA strand discontinuities in transcriptional regulation and in higher-order chromatin organization are integrated.

## Acknowledgements

The authors thank Kristian Helin and Karl Agger (Copenhagen, Denmark) for the KDM4 KO mES experimental system, Hiroshi Kimura (Tokyo, Japan) for the histone PTM-specific antuibody panel and György Fenyőfalvi for performing the halo winding assay.

## Materials and Methods

### Cells

HeLa, H3-GFP and H4-GFP expressor HeLa ^83^ (provided by Hiroshi Kimura,), were cultured in DMEM supplemented with 10% FCS, 2 mM L-glutamine, 100 μg/ml streptomycin, 100 U/ml penicillin. Jurkat and NALM6 cells were cultured in RPMI supplemented with 10% FCS, 2 mM L-glutamine, 100 μg/ml streptomycin, 100 U/ml penicillin. mES (from Invitrogen) and mES KDM4 KO cells (provided by Kristian Helin) were cultured and treated with 4-OHT as described in ^54^. NALM6 and SY-5Y cells (from C. A. Austin’s lab) were cultured in MEM/F12 supplemented with 10% FCS, 2 mM L-glutamine, 100 μg/ml streptomycin, 100 U/ml penicillin.

### Embedding live cells into low melting point agarose

Prior to embedding, the wells of 8-well chambers (Ibidi, Martinsried, Germany) were coated with 1% (m/v) low melting point (LMP) agarose. 150 μl liquid agarose, diluted in distilled water was dispensed into each well and was immediately removed so that a thin agarose layer remained on the surfaces and was left to polymerize on ice for 2 minutes, then kept at 37°C until the surface of the wells dried out. This coating procedure was repeated once more on the same chambers. Embedding was performed keeping cells and agarose at 37°C. The cell suspension containing 6×10^6^ cells/ml was mixed with 1% LMP agarose diluted in 1×PBS (150 mM NaCl, 3.3 mM KCl, 8.6 mM Na2HPO4, 1.69 mM KH2PO4, pH 7.4) at a v/v ratio of 1:3. 22 μl of the cell-agarose suspension was dispensed in the middle of the wells and the chambers were covered with home-made rectangular plastic coverslips cut out from a 200 μm thick medium weight polyvinyl chloride binding cover (Fellowes, Inc., Itasca, Illinois, USA). Cells were left to sediment on the surface of the coated wells for 4 minutes at 37°C, then kept on ice for 2 minutes. After polymerization of the agarose, 300 μl ice cold complete culture medium was added to each well, a step aiding removal of the coverslips and used for cell treatment (below) at the same time.

### Histone eviction by intercalators

The agarose embedded cells at the bottom of the wells were washed with 500 μl ice cold 1×PBS, three times for three minutes each, then permeabilized with 500 μl ice cold 1% (v/v) Triton X-100 dissolved in 1×PBS/EDTA (5 mM EDTA in PBS), for 10 minutes. This step was repeated once more. After permeabilization, nuclei were washed with 500 μl ice cold 1×PBS/EDTA three times for three minutes and were treated with different concentrations of intercalator solutions on ice. Ethidium bromide (EBr) (Thermo Fisher Scientific, Waltham, Massachusetts, USA) diluted in 1×PBS/EDTA and supplemented with 600 mM NaCl was added to the final concentrations indicated in the figures. Doxorubicin (TEVA, Debrecen, Hungary) was diluted in 1×PBS/EDTA when added to the permeabilized nuclei. Nuclei were incubated with 500 μl of ice cold intercalator solution for 60 (EBr) or 120 (Doxorubicin) minutes. After this treatment, nuclei were washed with 500 μl ice cold 1×PBS/EDTA three times for 3 minutes. Analysis of the curves was made by SigmaPlot 12.0, using either ‘Sigmoid 3 parameter’ (in the case of linear plots) or ‘Standard curves: Four Parameter Logistic Curve’ (in the case of logarithmic plots) curve-fitting subroutines. Elution curves were normalized to ’1’ dividing the mean fluorescence intensities represented by the data points by that of the non-treated sample. All the SEM values indicated in the Figures were calculated from the datapoints of the cell population analyzed in the given experiment.

### Immunofluorescence labeling

After permeabilization and intercalator or enzymatic treatment the samples were incubated with 500 μl 5% (m/v) Blotto Non-Fat Dry Milk (Santa Cruz Biotechnology Inc., Santa Cruz, California, USA) in 1×PBS/EDTA for 30 minutes on ice to decrease nonspecific binding of the antibodies. The blocking solution was washed out with 500 μl ice cold 1×PBS/EDTA three times for three minutes and indirect immunofluorescence labeling was performed using mouse monoclonal anti-H3K4me0/1/2/3 (^84^; stock: 0.5 mg/ml), mouse monoclonal anti-H3K27me1/2/3 (^84^; stock: 0.5 mg/ml), mouse monoclonal anti-H3K9me1/2/3 (^84^; stock: 0.5 mg/ml), mouse monoclonal anti-H3K9ac/14ac/27ac (^84^; stock: 0.5 mg/ml) and mouse monoclonal anti-H4K8ac/16ac (^84^; stock: 0.5 mg/ml) primary antibodies, all diluted in 150 μl of 1×PBS/EDTA/1% bovine serum albumin (BSA); (1×PBS/EDTA supplemented with 1% w/v BSA), at 4°C, overnight. All the above antibodies were applied to the wells at a titer of 1:800. After labeling with the primary antibodies, the nuclei were washed with 500 μl ice cold 1×PBS/EDTA three times for 10 minutes. Labeling with the secondary antibodies was performed in 150 μl 1×PBS/EDTA for two hours on ice using Alexa fluor 488 (A488) or Alexa fluor 647 (A647) conjugated goat anti-mouse IgG or goat anti-rabbit IgG antibodies (Thermo Fisher Scientific, Waltham, Massachusetts, USA). The secondary antibodies were also used at a titer of 1:800, diluted in 1×PBS/EDTA, prepared from 2 mg/ml stock solutions. After labeling with the secondary antibodies, the agarose embedded nuclei were washed with 500 μl ice cold 1×PBS/EDTA three times, for 10 minutes. Then the samples were fixed in 1% paraformaldehyde (dissolved in 1×PBS/EDTA) at 4°C, overnight. After fixation the wells containing the embedded nuclei were washed with 500 μl ice cold 1×PBS/EDTA three times, for 3 minutes and were stained with 200 μl 25 μg/ml propidium–iodide (PI, dissolved in 1×PBS/EDTA) for 30 minutes, on ice. The stained nuclei were washed three times with 500 μl ice cold 1×PBS/EDTA for 3 minutes. Fluorescence intensity distributions were recorded using an iCys laser scanning cytometer (LSC).

### Etoposide treatment

Mouse ES cells were treated with etoposide (TEVA, Debrecen, Hungary) at a final concentration of 250 µM. The drug was diluted in complete DMEM medium and the cells were incubated together with the drug at 37°C in 5% CO2 atmosphere, for 1 hr. Agarose embedded Jurkat cells were treated with 20 µM and 200 µM etoposide dissolved in RPMI for 30 minutes at 37°C, in a 8-well Ibidi slide.

### Nickase and DNase I treatment

The frequent cutter Nt.CviPII nickase (recognition site: CCD; New England Biolabs Inc., Ipswich, Massachusetts, USA; 5000 U/ml) and DNase I (2 mg/ml) enzymes were applied after the washing steps following permeabilization (see above). Before digestion, agarose was equilibrated with nickase buffer (10 mM Tris-HCl pH 8, 50 mM NaCl, 10 mM MgCl2, 1 mg/ml BSA) or with DNase I buffer (10 mM Tris-HCl pH 8, 0.1 mM CaCl2, 2.5 mM MgCl2) by washing three times with 500 µl buffer solutions. Nickase treatment was performed in 300 µl nickase buffer for 30 min at 37 °C, using the enzyme at a final concentration of 0.5 U/ml. DNase I digestion was performed in 300 µl DNase I buffer for 10 min at 37 °C, at a final concentration indicated in the figures. After enzymatic treatments, the samples were washed with 500 μl ice cold 1×PBS/EDTA three times for three minutes.

### Automated microscopy

Automated microscopic imaging was performed using an iCys instrument (iCys® Research Imaging Cytometer; CompuCyte, Westwood, Massachusetts, USA). Green fluorescent protein (GFP), SYBR Gold, A488, doxorubicin and PI were excited using a 488 nm Argon ion laser, A647 with a 633 nm HeNe laser. The fluorescence signals were collected via an UPlan FI 20× (NA 0.5) objective. GFP and A488 were detected through 510/21 nm and 530/30 nm filters, respectively, while doxorubicin, A647 and PI were detected through a 650/LP nm filter. Each field (comprising 1000×768 pixels) was scanned with a step size of 1.5 µm. In the case of the winding assay SYBR Gold fluorescence was collected via a 10x objective and detected through a 550/30 nm filter. Amplification of the SYBR Gold signal via PMT voltage and gain were adjusted for each well separately so that the matrix area of the G1 phase halos (at a threshold value of 4000 pixel intensity) were kept constant. Background subtraction (offset) was adjusted so that the background pixel intensity was set around 300 in each well. The average halo radii for G1 cells were calculated from the area of the halos (measured at a threshold value of 600 pixel intensity). Data evaluation and hardware control were performed with the iCys 7.0 software for Windows XP. Gating of G1 phase cells was according to the fluorescence intensity distribution of the DNA labeled with PI.

### Genome-wide mapping of chromosomal nicks (NLDI)

Lymphocytes were isolated from human peripheral blood on Ficoll, fixed in 1% paraformaldehyde (30 mins, RT) and the excess formaldehyde was quenched with 0,7 M glycine. 90 µl agarose plugs were prepared according to standard protocols. The agarose encapsulated cells were lysed and deproteinized in the following buffer: 0,43 M EDTA, 1 v/v% Sarcosyl, 0,01 M Tris (pH=8) and 15 unit/ml Proteinase K. The blocks were incubated at 55 °C for 48 hours and the lysing/deproteinizing buffer was changed for a fresh aliquot after 24 hours. Nick tagging was performed in the presence of chain terminator ddNTPs by limited *in situ* nick translation using the E coli DNA polymerase I holoenzyme (DNA POL I). The agarose plugs were equilibrated in 2 ml Eppendorf tubes for 3×50 minutes in DNA POL I buffer and then transferred to ice for 30 minutes in the nick translating mixture consisting of 1x DNA POL I buffer, 5 µM of each ddNTPs, 1 µM of dATP, dGTP, biotin-dCTP, biotin-dUTP, and 150 units/ml of DNA POL I. Nick tagging was initiated by transferring the tubes to 37 °C for 30 minutes with gentle shaking. The reaction was stopped by washing the agarose plugs in excessive amounts of 0.5 M EDTA. DNA immunoprecipitation was carried out as follows: 4-6 agarose plugs were digested with ß-agarase (2 units/plug), sonicated for 15’ at „high” setting with a Diagenode Bioruptor and then purified using a Macherey-Nagel PCR cleanup kit (according to manufacturer’s protocol). Immunoprecipitation of nicks were carried out according to standard ChIP protocols using Protein G-coupled Dynabeads coated with monoclonal anti-biotin antibodies (Sigma-Aldrich) antibody. Libraries were prepared according to the Illumina’s standardized protocol. Each library was denatured by 1N NaOH and diluted to 10 pM final concentration. Cluster generation was performed on cBot instrument using TruSeq SR Cluster Kit v3 according to manufacturer’s protocol. Each library pool was loaded into 3 - 3 lanes of the flow cell, then single read 50 bp sequencing runs were performed on an Illumina HiScan SQ instrument (Illumina) using TruSeq SBS Kit v3 - HS (50-cycles). Library preparations and sequencing were executed by Genomic Medicine and Bioinformatic Core Facility of University of Debrecen, Hungary.

### Nuclear halo preparation for immunofluorescence staining of nicks, topoisomerases and Lamin B

The cells were washed in PBS and resuspended in pre-warmed (37°C) 1xPBS to a density of 6 million/ml and embedded into LMP agarose in the wells of an Ibidi chanber. After the gel has set, the cells were washed three times in 1×PBS (500 µl/well) on ice. Lysis was performed using a Lysis Buffer consisting of 0.5 v/v% TritonX-100, 300 mM NaCl, 20 mM tris, 5 mM EDTA, 1 mM EGTA (400 µl/well, 10 minutes at room temperature (RT)). A high salt elution buffer (2 M NaCl, 20 mM TRIS-HCl, 5 mM EDTA, 1 mM EGTA) was applied to wash out most DNA-associated proteins (400 µl/well, 10 minutes at RT). Topoisomerases were stained using TOP2α or TOP2β specific antibodies (from Santa Cruz or C. A. Austin’s lab, at 1:100 dilution in 5 mg/ml BSA/PBS/EDTA, 4°C, overnight). Lamin B was detected using anti-Lamin B antibody produced in goat and Alexa488 conjugated donkey anti-goat seconday antibody (1:200 dilution in 5 mg/ml BSA/PBS/EDTA, 4°C, overnight). Imaging was carried out using an Olympus FluoView 1000 confocal laser scanning microscope (60x oil immersion objective, 488nm and 633 lasers).

### Confocal microscopy

Imaging was carried out in an Olympus FluoView 1000 CLSM equipped with 488 and 633 nm lasers, using a 60x oil immersion oil objective or in a Nikon N-STORM Ti-E inverted confocal microscope, with a 60x water immersion objective. Composite images were constructed and evaluated using the ImageJ software. Instrument settings (laser power, PTM voltage, gains, picture dwell time) and image analyses parameters (brightness, contrast, gamma factor of ImageJ) were identical in the case of all the samples compared in a particular experiment.

### Preparation of agarose plugs containing genomic DNA

Preparation of agarose-plugs was carried out by the standard methods. Briefly, cells were harvested and washed twice in PBS/EDTA. Samples were mixed with an equal volume of 1.5% low melting point (LMP) agarose (Sigma-Aldrich) dissolved in PBS/EDTA. Aliquots were allowed to harden in sample molds at 4°C for 5 min. Each plug contained ∼2.5 × 10^6^ cells. The plugs were digested with 0.5 mg/ml Proteinase K (Thermo Fisher Scientific) in lysing solution (0.5 M EDTA, 10 mM Tris–HCl, 1% Sodium lauroyl sarcosinate (Sarkosyl), pH 8.0) at 55°C for 2 days, then washed with TE (10 mM Tris–HCl, 2 mM EDTA, pH 8.0) and treated by 0.75 μM phenyl-methyl-sulfonyl-fluoride (PMSF, Sigma-Aldrich) at 37°C for 10 min in order to inactivate residual proteinase activity. Finally, the plugs were washed with TE and stored in the same buffer at 4°C.

### Molecular combing

Molecular combing was performed on the genomic DNA samples of mES KDM4 KO cells as described in ref. ^55^. Genomic DNA embedded in agarose plugs prepared and nick-labeled using biotinylated nucleotides as described above. To solubilize the DNA, 1.6 ml 0.1 M MES (pH 6.5) was added to each plug, incubated at 70°C for 20 min, then at 42°C for 10 min. The blocks were dissolved by treatment with 8 U Agarase (Thermo Fisher Scientific; at 42°C overnight). The DNA solutions were placed at RT and transferred to disposable reservoirs without pipetting. DNA combing was performed by the combing apparatus of Genomic Vision (France) according to the manufacturer’s instructions using 22×22 mm vinylsilane coated coverslips obtained from the same source. After combing, the coverslips were glued to glass slides with cyanoacrylate. Non-specific binding of the antibodies was blocked by incubation with 30 µl of 5% BSA/1×PBS/0.1% Triton X-100 for 20 min in a humid chamber. The biotin molecules incorporated into the DNA were visualized by indirect immunofluorescent labeling using 1:60 diluted mouse anti-biotin as a primary antibody (Sigma-Aldrich) applied in 1% BSA/1×PBS/0.1% Triton X-100 for 45 min at RT in a humid chamber. After washing 3 times with 3 ml 1×PBS, twice with 30 µl PBS/0.1% Triton X-100 and then once with 30 µl PBS for 5 min, Alexa Fluor 647 conjugated goat anti-mouse antibody (Life Technologies) was used as a secondary antibody at a final concentration of 17 µg/ml in 1% BSA/1×PBS/0.1% Triton X-100, at RT in a humid chamber, for 45 min. Following labeling the coverslips were washed as before. DNA staining was performed using 30 µl YOYO-1 dye (Thermo Fisher Scientific) diluted 1:5000 in 0.1 M MES (pH 6.5) in a humid chamber, in the dark, for 20 min. The coverslips were covered by ProLong® Gold Antifade using a clean coverslip and placed at 4°C overnight. Imaging was carried out using a Nikon N-STORM Ti-E confocal laser scanning microscope equipped with 488 and 633 nm lasers, using a 60× water objective. The pictures were evaluated by ImageJ, graphs were plotted by Sigma Plot 11.0 software.

### Gel electrophoretic and reverse Southwestern blot (rSW) analyses

Standard, nondenaturing agarose gelelectrophoresis was perfomed with or without posttreatment of the nickase treated agarose-embedded H2B-GFP expressing samples with S1 nuclease (1000 U/ml) in its own buffer (1h at 37°C), to convert nicks into double-strand breaks. For pulse-field gelectrophoresis of the S1- treated samples a CHEF mapper XA Pulse Field Electrophoresis System (Bio-Rad Laboratories Inc., Hercules, California, USA) was used following the manufacturer’s instructions to resolve 3-300 kb fragments. For size markers O’GeneRuler 1kb DNA ladder (Thermo Fisher Scientific, Waltham, Massachusetts, USA), lambda DNA (Thermo Fisher Scientific, Waltham, Massachusetts, USA) and Pulse Marker 2200-225 kb were used.

Pulse-field gel electrophoresis of agarose embedded samples was performed by a CHEF mapper XA Pulse Field Electrophoresis System (Bio-Rad Laboratories Inc., Hercules, California, USA) (1% agarose gel (in 0.5% TBE); automatic program; separation range: 5-150 kb). As DNA ladders MidRange PFG Marker (New England Biolabs, UK) was used. The samples were digested with the rare-cutting restriction endonuclease SfiI at 20 U/sample (Thermo Fisher Scientific, Waltham, Massachusetts, USA) prior to electrophoresis in order to increase DNA fragmentation. The gel was stained with 5 µg/ml EBr. rSW blotting was carried out following a protocol developed earlier ^55^. Briefly, gels were vacuum-blotted on Immobilon P Transfer Membrane (PVDF, 0.45 µm, Millipore) and cross-linked to the membrane by UVC. The membranes were pre-hybridized for 1 hour at RT in 15 ml pre-hybridization solution (5% BSA, 0.2% Tween-20/1×PBS), then incubated with mouse anti-biotin primary antibody at a dilution of 1:2000 in 15 ml hybridization solution for 16 hours at 4 °C. After washing with 0.2% Tween-20/1×PBS for 5×5 min, the membranes were incubated with goat anti-mouse IgG antibody conjugated with horseradish peroxidase (dilution 1:2500) at RT, for 1.5 h. The chemiluminescence signals were detected by a ChemiDoc Touch Imaging System (Bio-Rad, Hercules, CA, USA). Blots were evaluated by the ImageJ software. For quantification, the biotin signal intensities of the fragments (in the size range of 30-150 kb) relative to their EBr signal intensities were calculated. The calculated biotin/EBr ratios of OHT treated or mutant cells were normalized to the ratios of untreated or control cells. Since the ratios are proportional to the incidence of nicks in a given amount of DNA (EBr signal) the observed changes are independent of differential loading.

### Nuclear halo preparation in 96 well plates

96 well tissue culture plates (TPP, Switzerland) were coated with α-poly-L-lysine (MW 150 000-300 000, Sigma-Aldrich, St-Louis, MO) by pipetting 100 μL polylysine solution (0.5 mg/mL, dissolved in water) into each well and incubating at RT for 2 h. After the removal of the solution the plates were dried at RT in a sterile hood and stored at RT (not any longer than a few days). Wells were washed 3⨉ with 200 μL H2O immediately before use, to rinse away unbound poly-lysine. All the following operations were carried out on ice. 100 μL aliquots of the nuclear suspension were added into the wells of the polylysine coated plates (∼1000 nuclei/well) and the nuclei were spun down to the bottom of the wells (2250 g, 10 min, 4°C) in a plate centrifuge. The supernatant was removed carefully using vacuum and 100 μL aliquots of the twisting solution series with the appropriate SYBR Gold concentration (see below) were applied immediately. The plates containing the nuclear halos at their bottom could be stored overnight in the dark at 4°C without any deterioration, prior to LSC analysis. SYBR® Gold (SG) nucleic acid gel stain was purchased from Life Technologies (Carlsbad, CA, USA). The dye is supplied at 10,000⨉ concentration in dimethyl sulfoxide (DMSO). Solutions of SG were prepared in nuclear extraction buffer (TRIS 20 mM, pH 7.5, NaCl 2 M, sucrose 250 mM, EDTA 5 mM, EGTA 1 mM). Since SYBR dyes bind to plastic surfaces to some extent, the plastic tubes used for the preparation of the dilution series were incubated overnight together with 1X SYBR Gold solution to saturate their surfaces with the dye and were rinsed 3X with H2O prior to their use. During the serial dilution procedure the same pipette was used for pre-dispensing the appropriate amount of diluent as well as for the consecutive transfers of the dye between the tubes, thus ensuring constant ratio between adjacent dilution steps.

### LSC analysis of nuclear halos

Plates containing the nuclear halos were allowed to warm up to RT before scanning. Halos were then analyzed using an LSC apparatus (specified above). Fluorescence was excited with a 488 nm argon laser, and emission was detected through a 550 (± 30) nm bandpath filter. The wells were scanned using a 20x objective. PMT voltage and gain were adjusted so that the intensity values of the brightest pixels of G2- and G1 phase halos were around 80% and 60% of the maximum intensity, respectively. The background subtraction (offset) was fine-tuned until the intensity of background pixels was around 300 (i.e. ∼2% of the dynamic range of the measurement). The average halo radii at different intercalator concentrations were calculated from the measured round halo areas and were plotted against the dye concentration, thus generating the characteristic ‘winding curve’.

