## Supplementary figures for "KDM4-dependent DNA breaks at active promoters facilitate +1 nucleosome eviction"

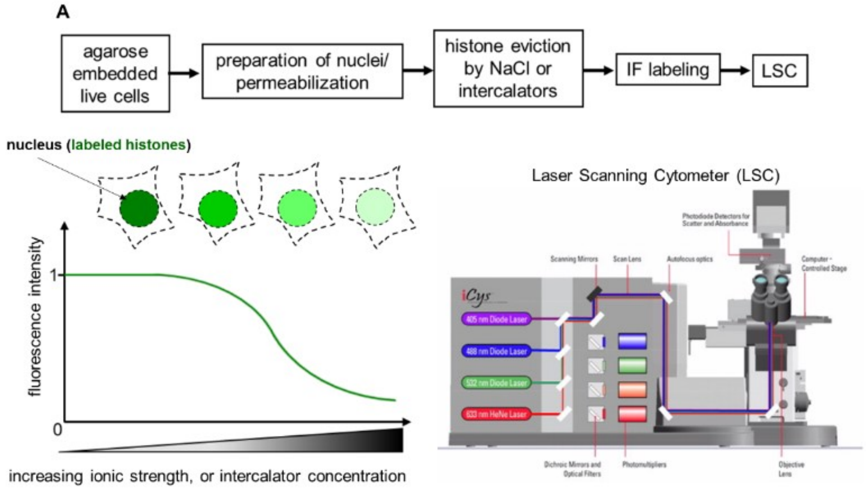


Suppl. Fig. 1

Flow-chart of the QINESIn assay (Quantitative Imaging of Nuclei after Elution with Salt/Intercalators assay; [1]). The intercalator elution format was used in the current study, since the sensitivity of the tetrasomes (carrying the PTMs distinguishing the +1 nucleosomes) to changes of superhelicity is detectable via intercalator, rather than salt, elution (as described in[1]). The histones remaining in the nuclei after treatment with increasing concentration of intercalator (ethidium bromide; EBr) solutions are detected by indirect immunofluorescence labeling (or directly if the histones are tagged with a fluorescent protein), and quantitatively analyzed by laser scanning cytometry (LSC) on a cell-by-cell basis. The LSC instrument used in the experiments is equipped with four lasers: 405 nm, 488 nm, 563nm, 633 nm and four photomultiplier tubes (PMTs), each detecting a specific wavelength range of fluorescence excited by the scanning lasers. As the laser light intersects the sample, scattered or transmitted light is also simultaneously directed to one or more photomultiplier tubes. The PMT signals are converted into images and the events, such as cells, nuclei or other subcellular structures are identified and segmented on the basis of their fluorescence. From the segmented events, a variety of quantitative data are calculated: area, integral fluorescence, maximal pixel intensity, circularity, perimeter, *x* and *y* coordinates, among others. The numerical values are displayed in scattergrams and histograms, allowing assessment of relationships among the various features.

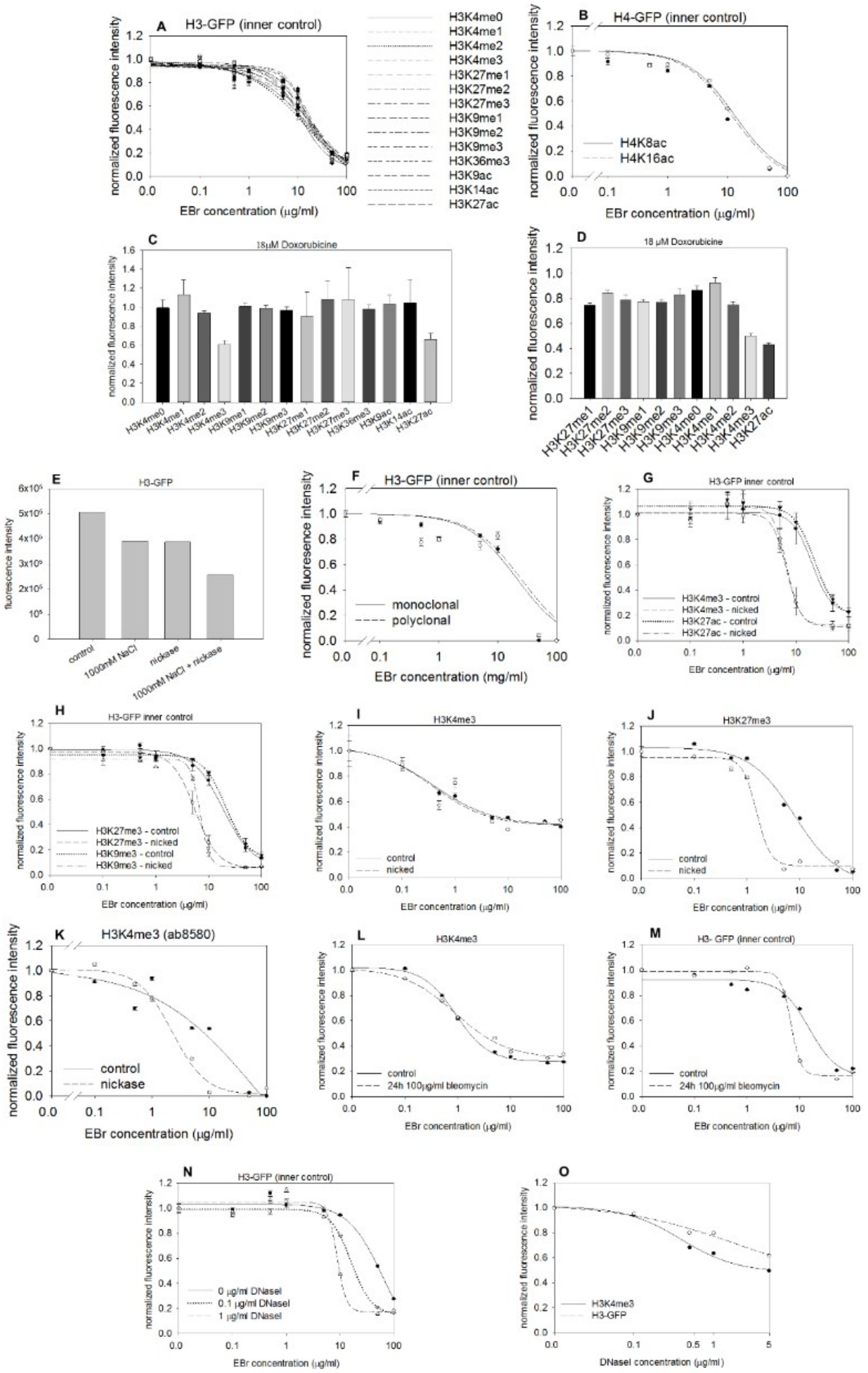
A
H3-GFP (inner control)
normalized fluorescence intensity
EBr concentration (µg/ml)
H3K4me0
H3K4me1
H3K4me2
H3K4me3
H3K27me1
H3K27me2
H3K27me3
H3K9me1
H3K9me2
H3K9me3
H3K36me3
H3K9ac
H3K14ac
H3K27ac
B
H4-GFP (inner control)
normalized fluorescence intensity
EBr concentration (µg/ml)
H4K8ac
H4K16ac
C
18µM Doxorubicine
normalized fluorescence intensity
H3K4me0
H3K4me1
H3K4me2
H3K4me3
H3K9me1
H3K9me2
H3K9me3
H3K27me1
H3K27me2
H3K27me3
H3K36me3
H3K9ac
H3K14ac
H3K27ac
D
18 µM Doxorubicine
normalized fluorescence intensity
H3K27me1
H3K27me2
H3K27me3
H3K9me1
H3K9me2
H3K9me3
H3K4me0
H3K4me1
H3K4me2
H3K4me3
H3K27ac
E
H3-GFP
fluorescence intensity
control
1000mM NaCl
nickase
1000mM NaCl + nickase
F
H3-GFP (inner control)
normalized fluoresence intensity
monoclonal
polyclonal
EBr concentration (mg/ml)
G
H3-GFP inner control
normalized fluorescence intensity
H3K4me3 - control
H3K4me3 - nicked
H3K27ac - control
H3K27ac - nicked
EBr concentration (µg/ml)
H
H3-GFP inner control
normalized fluorescence intensity
H3K27me3 - control
H3K27me3 - nicked
H3K9me3 - control
H3K9me3 - nicked
EBr concentration (µg/ml)
I
H3K4me3
normalized fluorescence intensity
control
nicked
EBr concentration (µg/ml)
J
H3K27me3
normalized fluorescence intensity
control
nicked
EBr concentration (µg/ml)
K
H3K4me3 (ab8580)
normalized fluorescence intensity
control
nickase
EBr concentration (µg/ml)
L
H3K4me3
normalized fluorescence intensity
control
24h 100µg/ml bleomycin
EBr concentration (µg/ml)
M
H3- GFP (inner control)
normalized fluorescence intensity
control
24h 100µg/ml bleomycin
EBr concentration (µg/ml)
N
H3-GFP (inner control)
normalized fluorescence intensity
0 µg/ml DNaseI
0.1 µg/ml DNaseI
1 µg/ml DNaseI
EBr concentration (µg/ml)
O
normalized fluorescence intensity
H3K4me3
H3-GFP
DNaseI concentration (µg/ml)

Suppl. Fig. 2.

QINESIn experiments using EBr elution.

(A and B) EBr elution curves of H3-GFP (A) and H4-GFP (B), used as inner control in the experiment shown in Fig. 1A-E. (C) Doxorubicin elution measured at 18 μg/ml concentration of the intercalator in HeLa nuclei. (D) Doxorubicin elution measured at 18 μg/ml concentration of the intercalator in hPBMCs. (E) Initial H3-GFP fluorescence intensities measured in the absence of EBr in the experiment shown in Fig. 1G. (F) H3-GFP used as inner control in the experiment shown in Fig. 1H. (G and H) EBr elution of H3-GFP used as internal control in the experiments shown in Fig. 2A and B. (I and J) EBr elution curves measured in the case of relaxed (nickase) or superhelical (control) state of the genomic DNA of hPBMCs, using H3K4me3 or H3K27me3 specific antibodies. (K) EBr elution of H3K4me3 nucleosomes measured in HeLa nuclei treated with nickase. H3K4me3 was detected by the ab8580 polyclonal antibody. (L) EBr elution of H3K4me3 nucleosomes measured in the nuclei of H3-GFP expressing HeLa cells treated with bleomycin. (M) EBr elution of H3-GFP used as internal control in the experiment shown in panel (L). (N) EBr elution of H3-GFP used as internal control in the experiments shown in Fig. 2F. (O) Initial fluorescence intensities of H3K4me3 and of H3-GFP in the absence of EBr, plotted as a function of DNase I concentration in the experiment shown in Fig. 2F. The curves refer to 200-2000 G1 phase cells gated according to their DNA content. Bars represent SEM.

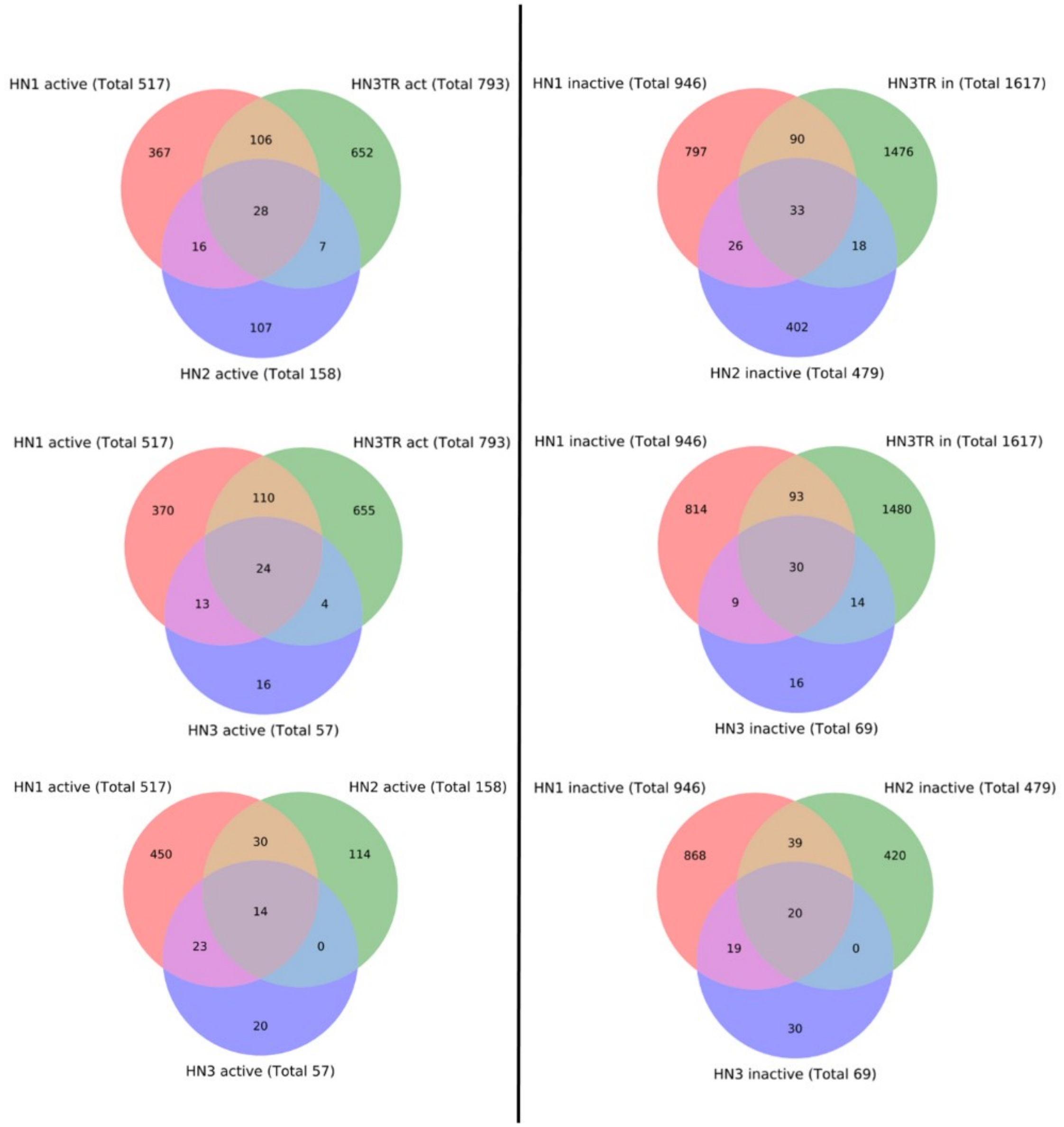


Suppl. Fig. 3.

Venn-diagram showing the comparison of active and inactive genes' lists containing nicked promoters in 4 experiments. In experiments labeled „HN1-3", nicks were labeled using DNA POL I, in „HN3TR" nicks were labeled using TdT and subsequently RNAse A-treated.

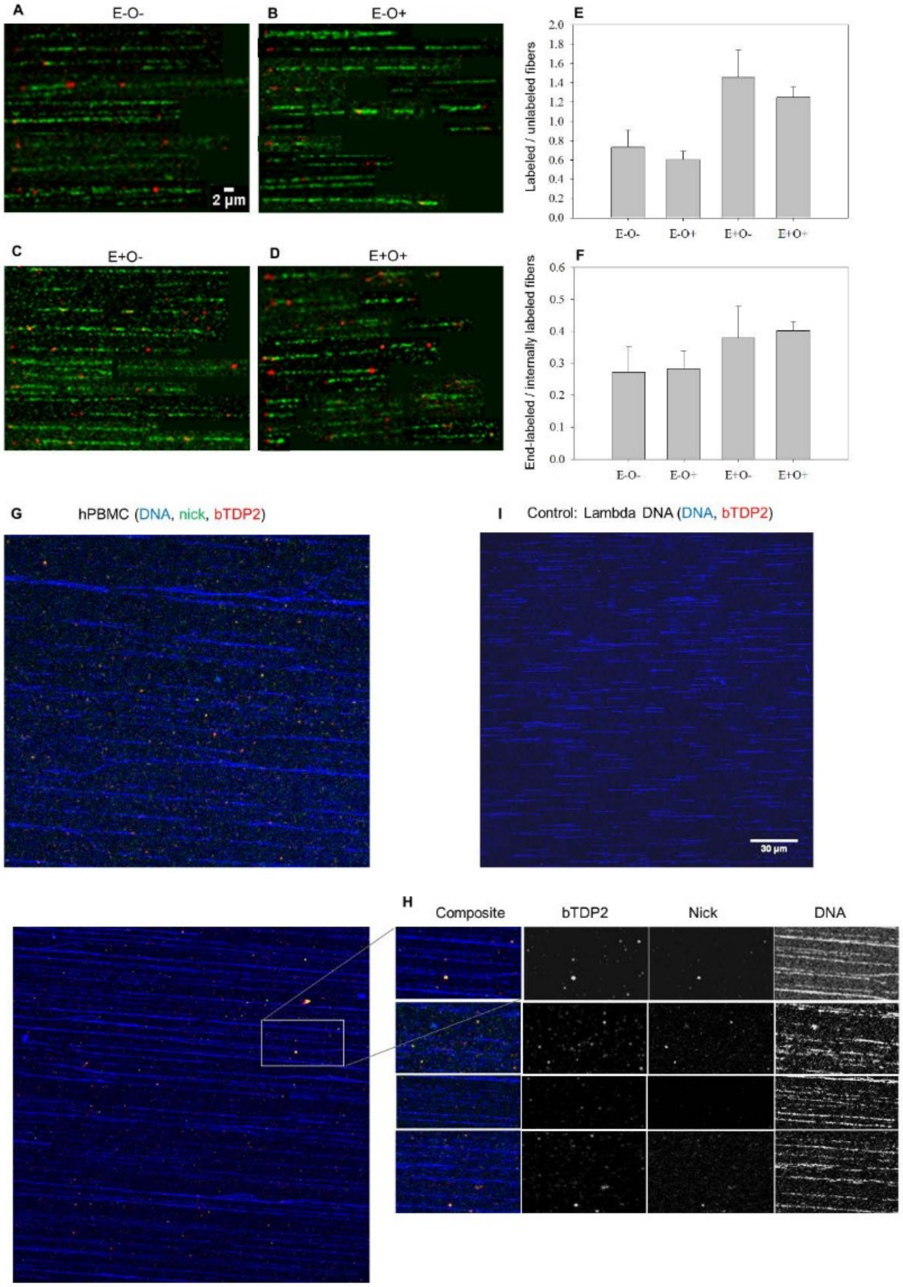


Suppl. Fig. 4.

(A-F) Molecular combing of nick-labeled genomic DNA of KDM4 KO mES control cells (panel A), of cells were treated with 4-OHT alone (panel B), treated with 250 µM etoposide alone (panel C), or co-treated with 4-OHT and etoposide (panel D). Biotinylated nucleotides were incorporated into agarose-embedded genomic DNA using DNA POL I. Biotin was detected by Alexa Fluor 647-conjugated anti-biotin antibody (red), the DNA molecules were stained with YOYO-1 (green). Panels A-D show mosaics of representative images cut out from larger field images. Panels E and F shows the quantitative analysis of labeling. Panel E: Ratio of labeled / unlabeled fibers (including fibers having more than one signal). Panel F: Ratio of end-labeled / internally labeled fibers. Fibers labeled on one or both ends were counted and divided by the number of fibers labeled internally. Fibers were counted in at least 3 vision fields. (G-H) Detection of endogenous 3'OH DNA breaks staining with biotin-labeled recombinant TDP2 (bTDP2). Nicks were labeled by incorporating ChromaTide 568 conjugated dUTP into hPBMC genomic DNA by DNA POL I (green). TOP2-related, DNA-bound phosphotyrosyl groups were detected by bTDP2, which was subsequently labeled by anti-biotin antibody (red). DNA was stained by YOYO-1 (blue). (G) DNA POL I and bTDP2 labeling in the case of hPBMC genomic DNA. (H) Composite and single-channel images (bTDP2, nick or DNA) showing a magnified part of the vision field in (G), and of 3 different vision fields. (I) Molecular combing of bTDP2 labeled Lambda phage DNA, used as control. Representative confocal microscopic images are shown in panels (G) and (I).

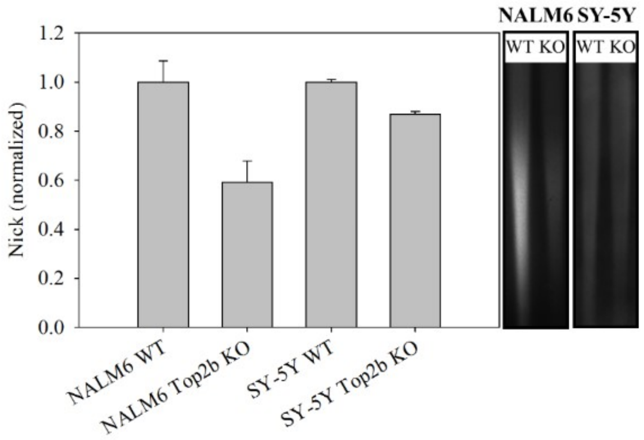


Suppl. Fig. 5.

rSW analyses of DNA POL I labeled nicks in TOP2β KO and wild type NALM6 and SY-5Y cells. The bar chart represents mean values ± SD obtained from three independent experiments for NALM6 and two independent experiments for SY-5Y samples. The inlet (on the right) shows the rSW anti-biotin staining of a blot of NALM6 WT, NALM6 TOP2β KO, SY-5Y WT, SY-5Y TOP2β KO cells in a representative experiment.

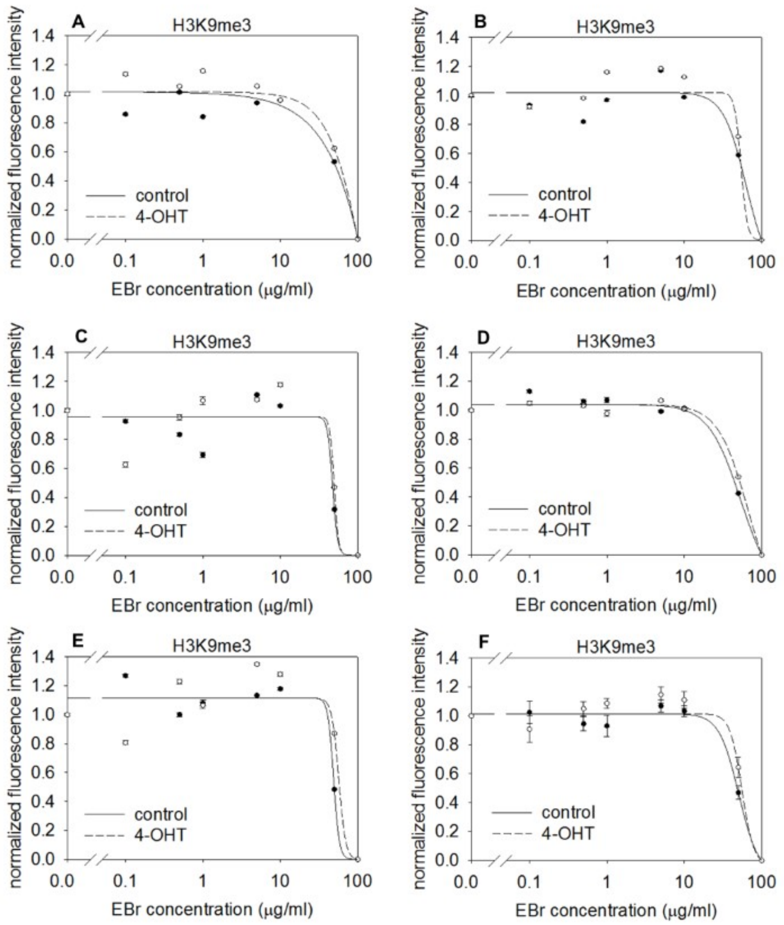


Suppl. Fig. 6.

(A-E) EBr elution of H3K9me3 nucleosomes measured in control and 4-OHT induced KDM4 KO mES nuclei. Results of five independent experiments are shown. H3K9me3 and H3K4me3 (data shown in Fig. 3A-E) nucleosomes were co-labeled. The curves refer to 200-2000 G1 phase cells gated according to their DNA content. Bars represent SEM. Average values and SEM of the five independent measurements are shown in panel (F).

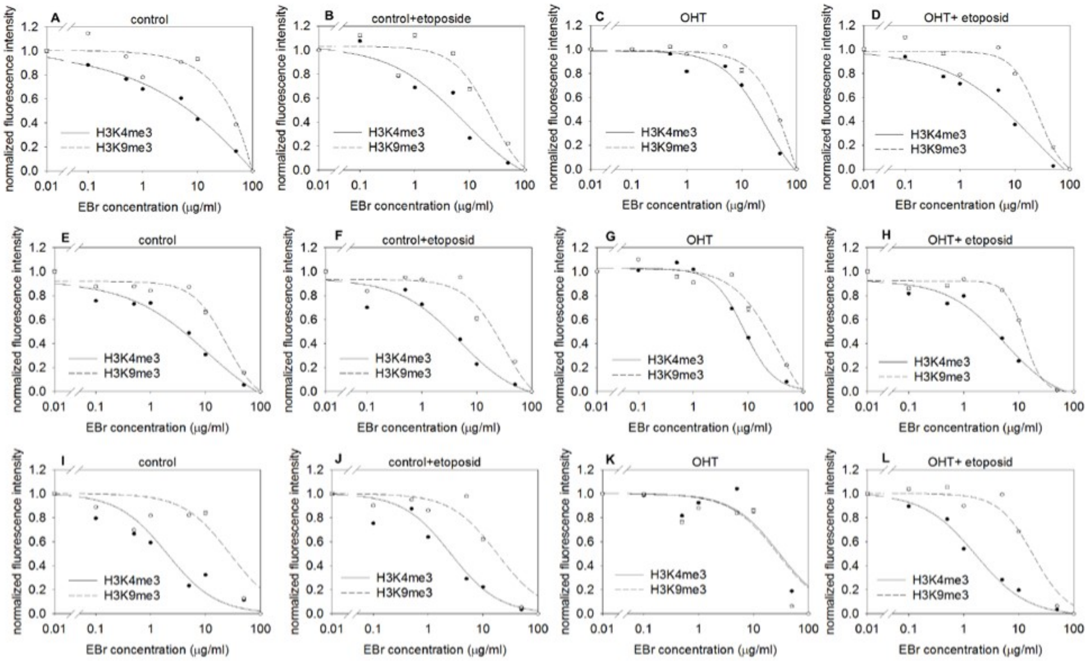


Suppl. Fig. 7.

(A-D) EBr elution of H3K4me3 nucleosomes measured in mES cells containing the KDM4 KO constructs, with (B and D) or without (A and C) etoposide treatment. Results of independent experiments are shown in panels (E-H) or on (I-L).

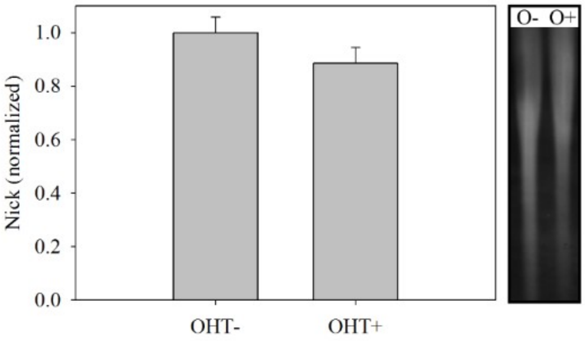


Suppl. Fig. 8.

rSW analyses of DNA POL I labeled nicks in KDM4 KO mES cells, with or without OHT treatment. Agarose embedded, deproteinized genomic DNA was nick-labeled and digested with SfiI. The fragments were separated by CHEF and transferred onto PVDF membrane. The incorporated biotin was labeled by anti-biotin antibody. Labeling intensities (biotin signal intensities of the fragments normalized to their corresponding EBr signal intensities) are shown in the 30-150 kb size range on the bar chart representing mean values ± SD obtained from eight independent experiments. The inset (on the right) shows the rSW anti-biotin staining of a blot in a representative experiment. See Materials and Methods for further details.

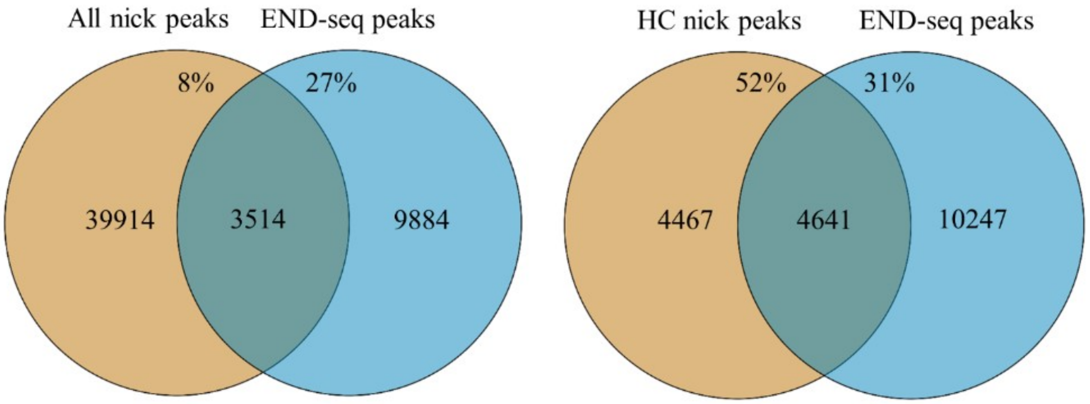


Suppl. Fig. 9.

Overlapping of DNA breaks detected by NLDI in hPBMCs with END-seq [2] peaks derived from NALM6 cells. The Venn-diagrams show the number of overlapping peaks. „HC nick peaks" marks a high confidence nick peakset (nick peaks present in at least two experiments). Percentages of overlapping peaks are also displayed. END-seq data were downloaded from the GEO database (GSM2635575, [3]).
